# Pleiotropic effects for Parkin and LRRK2 in leprosy type-1 reactions and Parkinson’s disease

**DOI:** 10.1101/507806

**Authors:** Vinicius M. Fava, Yong Zhong Xu, Guillaume Lettre, Nguyen Van Thuc, Marianna Orlova, Vu Hong Thai, Geison Cambri, Shao Tao, Ramanuj Lahiri, Linda Adams, Aurélie Cobat, Alexandre Alcaïs, Laurent Abel, Erwin Schurr

## Abstract

Type-1 reactions (T1Rs) are pathological inflammatory episodes and main contributors to nerve damage in leprosy. Here, we evaluate the gene-wise enrichment of rare protein altering variants in seven genes where common variants were previously associated with T1R. We selected 474 Vietnamese leprosy-patients of which 237 were T1R-affected and 237 were T1R-free matched controls. Gene-wise enrichment of nonsynonymous variants was tested with both kernel based (SKAT) and burden methods. Of the seven genes tested two showed statistical evidence of association with T1R. For the *LRRK2* gene an enrichment of nonsynonymous variants was observed in T1R-free controls (*p* SKAT-O= 1.6×10^−4^). This gene-wise association was driven almost entirely by the gain of function variant R1628P (*p* = 0.004; OR = 0.29). The second gene-wise association was found for the Parkin coding gene *PRKN* (formerly *PARK2*) where seven rare variants were enriched in T1R-affected cases (*p* SKAT-O = 7.4×10^−5^). Mutations in both *PRKN* and *LRRK2* are known causes of Parkinson’s Disease (PD). Hence, we evaluated to what extent such rare amino acid changes observed in T1R are shared with PD. We observed that nonsynonymous T1R-risk mutations in Parkin were enriched for amino acid mutations implicated in PD (*p* = 1.5×10^−4^). Hence, neuro-inflammation in PD and peripheral nerve damage due to inflammation in T1R share overlapping mechanisms of pathogenicity.

---

Leprosy is characterized by chronic infection with *Mycobacterium leprae* where the progression to disease is strongly dependent upon the host genetic background ^1–6^. One focus of current leprosy control efforts is the prevention of nerve damage ^7^. A major contributor to nerve damage are sudden-onset episodes of excessive inflammatory responses termed type-1 reactions (T1R) ^8^. Without timely intervention T1R can cause irreversible nerve damage due to a pathological cell-mediated inflammatory response directed against host peripheral nerve cells ^9^. Depending on the epidemiological setting, T1R episodes can afflict 30% to 50% of leprosy cases but are more common among older patients ^10^. While approximately one third of T1R are diagnosed at the time of leprosy diagnosis, T1R episodes can occur years after successful completion of therapy presumably in the absence of *M. leprae* bacilli ^10,11^.

Host genetic predisposition is an important aspect of T1R pathogenesis. A prospective study of host responses to *M. leprae* antigen identified a gene-set signature that predisposed to T1R ^12^. In addition, candidate gene approaches identified six protein coding genes in association with T1R ^13–19^. More recently a genome-wide association study (GWAS) identified a long noncoding RNA in chromosome 10 as T1R risk factor in independent populations and also provided suggestive evidence for the *PRKN* (formerly *PARK2*) as T1R susceptibility gene ^18^. Here, we tested if nonsynonymous variants with a likely functional impact at the protein level were enriched in the seven coding genes previously associated with T1R. We identified protein altering variants in the *PRKN* and *LRRK2* genes as T1R risk modulating factors. Since *PRKN* and *LRRK2* are established Parkinson’s disease (PD) susceptibility genes, we tested if T1R risk variants had been implicated in PD. Indeed, we found T1R risk modulating amino acid changes significantly enriched among PD susceptibility variants. Parkin loss-of-function mutations were shared risk factors for T1R and PD while a gain-of-function amino acid substitution in LRRK2 had an antagonist pleiotropic effect and was protective for T1R and a risk factor for PD.

## Results

We selected from our sample of Vietnamese leprosy patients 237 T1R-affected subjects (cases), which we matched by age at leprosy diagnosis, gender and clinical subtype of leprosy with 237 T1R-fee subjects as controls for deep re-sequencing of seven T1R susceptibility genes (Table S1, Table S2). We focused our analysis on the 63 variants that represented nonsynonymous changes, or insertion/deletions, and tested those for association with T1R. None of the studied subjects were homozygous or compound heterozygotes for nonsynonymous rare variants in any given gene. Of the seven genes, only non-synonymous changes in *PRKN* and *LRRK2* showed significant evidence for association with T1R (Table 1).

**Table 1.** Gene-wise association of protein altering variants in T1R-risk loci

| Gene | Var | Sing | No MAF cut off |  | Var | MAF < 0.05 |  |
| --- | --- | --- | --- | --- | --- | --- | --- |
| | | | $p_{\text{SKAT-O}}$ | $p_{\text{VT}}$ | | $p_{\text{SKAT-O}}$ | $p_{\text{VT}}$ |
| *TLR1* | 15 | 6 | 0.69 | 0.62 | 13 | 0.68 | 0.39 |
| *TLR2* | 9 | 7 | 0.11 | 0.47 | 9 | 0.11 | 0.45 |
| *PRKN* | 9 | 6 | $7.4 \times 10^{-5}$ | 0.01 | 7 | $1.1 \times 10^{-3}$ | $5.8 \times 10^{-3}$ |
| *TNFSF15* | 2 | 0 | 0.17 | 0.41 | 2 | 0.18 | 0.44 |
| *LRRK2* | 20 | 10 | $1.6 \times 10^{-4}$ | $2.2 \times 10^{-4}$ | 17 | $1.5 \times 10^{-4}$ | $1.4 \times 10^{-4}$ |
| *NOD2* | 10 | 5 | 0.61 | 0.97 | 10 | 0.62 | 0.98 |
| Combined | 65 | 34 | 0.04 | 0.34 | 58 | 0.04 | 0.37 |

Abbreviation: Var, number of nonsynonymous variants found for the gene; Sing, number of singletons variants found in only one individual; MAF, minor allele frequency.

### The LRRK2 R1628P variant is associated with protection from T1R

Nonsynonymous *LRRK2* variants were significantly enriched among T1R-free controls (Figure 1A, Table 1). Common variants did not contribute to this effect since restriction of the analysis to variants with MAF < 0.05 did not cause a decay in the strength of *LRRK2* association with T1R (Table 1). In contrast, the two low-frequency variants P755L and R1628P were highly enriched (>3-fold excess) in T1R-free subjects. When we performed single variant analyses we observed a strong protective effect of the LRRK2 R1628P mutation for T1R (*p* = 0.004, OR (95% CI) = 0.29 (0.13 - 0.67) for 1628P carriers). None of the other variants, including P755L, displayed significant univariate association with T1R (Table S3). We conditioned the gene-wise association of *LRRK2* with T1R on the R1628P mutation and observed a large drop in the strength of association (*p* SKAT-O = 1.6×10^−4^ to *p* SKAT-O = 0.01). By also adjusting on the P755L variant, *LRRK2* association with T1R protection became nominally non-significant (*p* SKAT-O = 0.06). However, given the remaining trend in favour of association it is possible that additional rare *LRRK2* variants contributed to protection from T1R.

**Fig 1.**
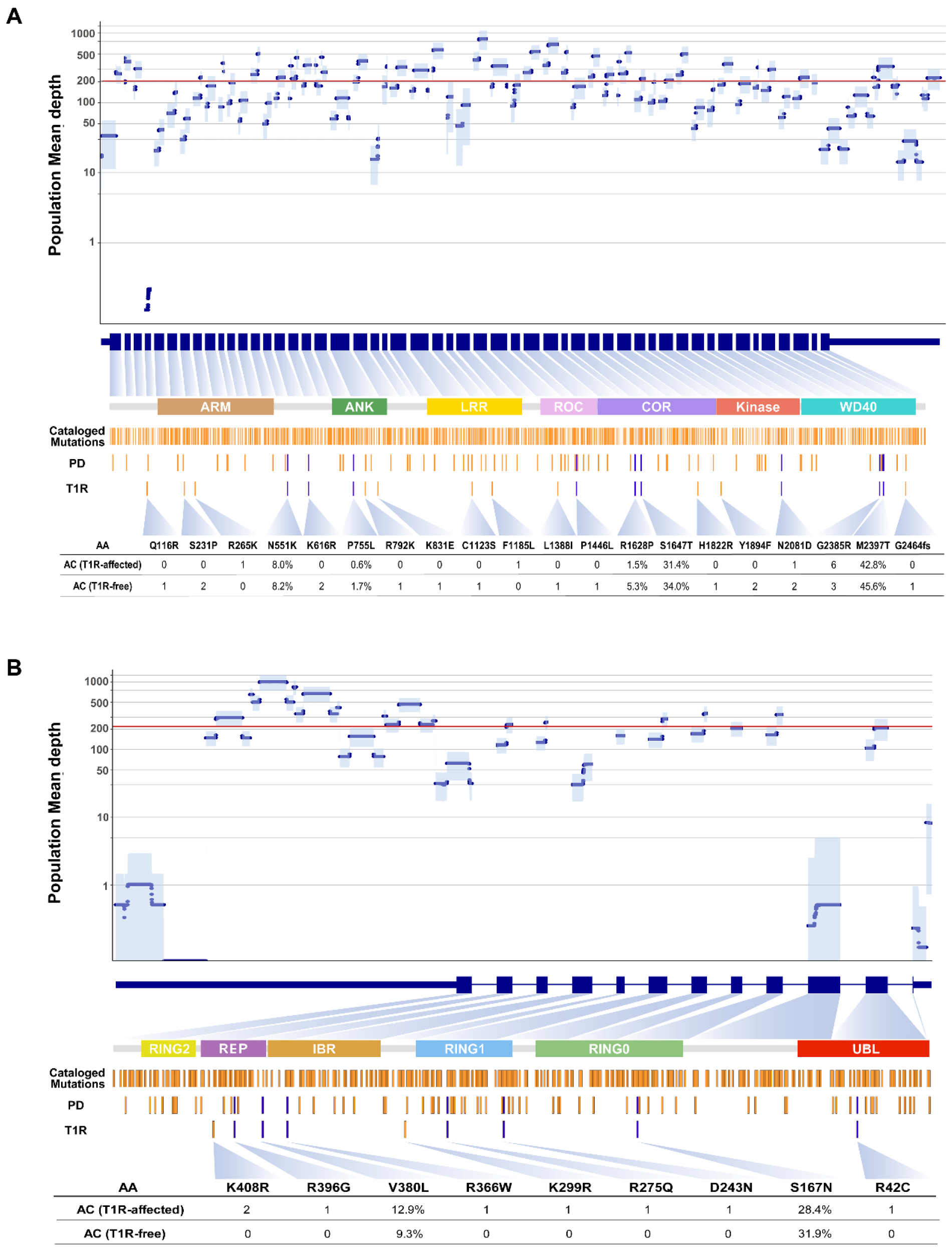
Overview of the deep-resequencing for the *LRRK2* and *PRKN* genes. At the top, the population mean depth of coverage is plotted according to the exons encoding the *LRRK2* gene (**A**) and *PRKN* (**B**). Blue circles indicate the depth of coverage per base-pair with the light blue shade representing the standard deviation of the mean. A red horizontal line marks the average depth of coverage for the two genes in the studied population. The protein functional domains are shown in the center and linked to their respective coding exons. The location of known coding mutation of LRRK2 and Parkin reported by either dbSNP or gnomAD, mutation associated with Parkinson’s Disease in PDmutDB and LOVD and mutations associated with Type-1 reaction are shown as yellow with blue bars denoting variants that overlap between PD and T1R. At the bottom, the allele count or the frequency of *LRRK2* and *PRKN* mutations are shown for the T1R-affected and T1R-free samples.

To validate the detected LRRK2 – T1R associations we decided to test for a possible impact of the 1628P and 755L mutations on LRRK2 functional activity. Hence, we genome-edited RAW cells to express hetero- and homozygously the 1628P and 755L mutant forms of *LRRK2*. We also generated a *LRRK2* knock-out line as negative control. Mutant and wildtype RAW cells were infected with *Mycobacterium bovis* (BCG) or *M. leprae*. Infection with BCG induced LRRK2 expression in all cells, but no significant difference of expression levels was observed between the LRRK2 wildtype and mutant proteins (Figure 2A, B). Following infection with BCG the 1628P mutant form mediated a significantly stronger respiratory burst while production of reactive oxygen species (ROS) by the 755L line was indistinguishable from the response of wildtype cells (Figure 2C, D). Compared to wildtype cells, the 1628P mutant also triggered a stronger respiratory burst in response to *M. leprae* (Figure 2E). Interestingly, the response of heterozygous LRRK2 1628 P/R cells was indistinguishable from the one mounted by homozygous 1628 P/P cells suggesting a dominant effect of the mutation (Figure 2F).

**Fig 2.**
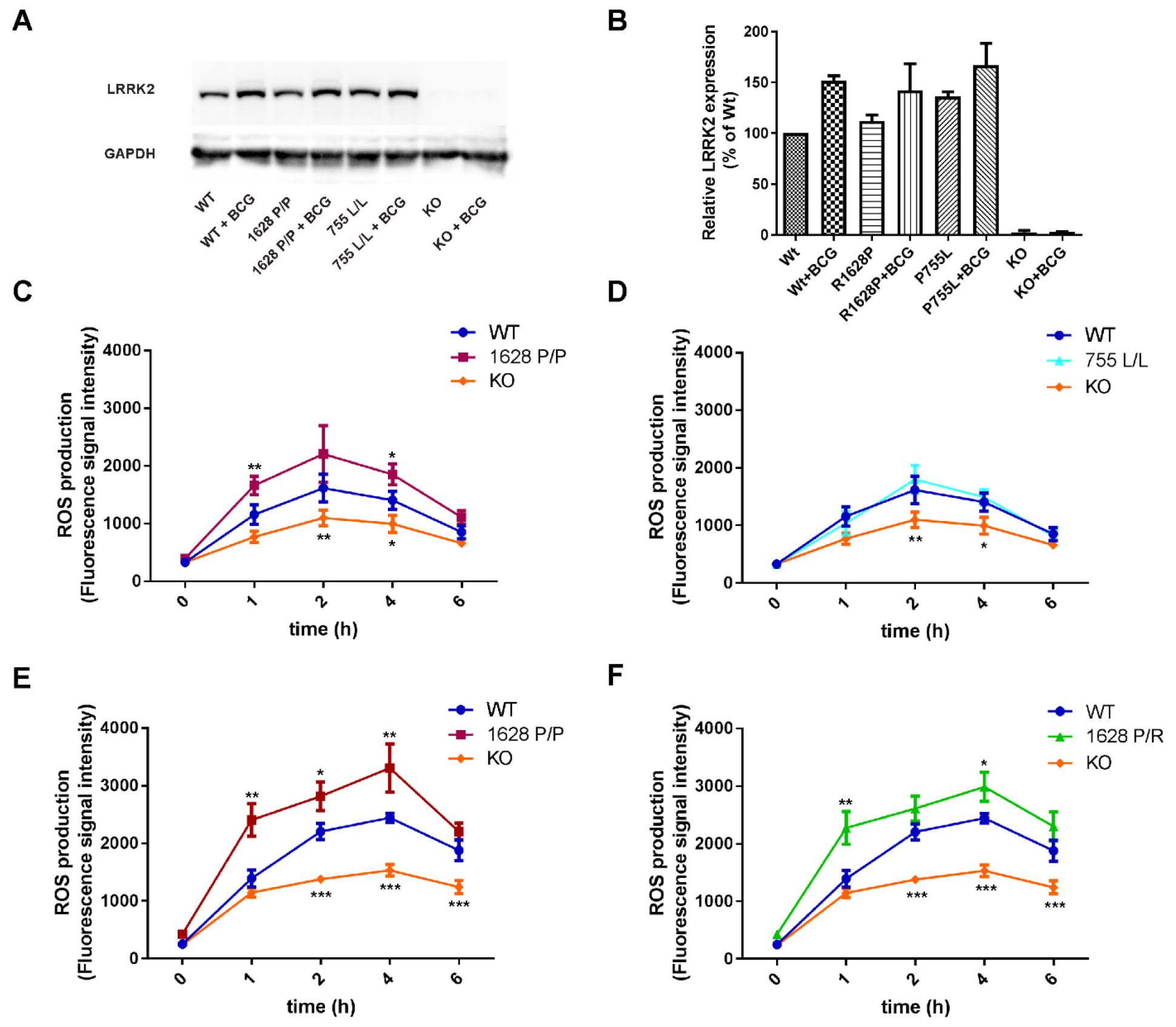
**The** LRRK2 R1628P rare variant displays an increased respiratory burst in response to challenge with both live BCG and *M. leprae*. Panels **A** and **B** show the relative abundance of LRRK2 for wild-type (WT) RAW cells and three CRISPR/Cas edited mutant cell lines in both resting state and under BCG stimulated conditions in a representative Western blot (**A**) and as triplicates with standard deviations (**B**). Panels **C** through **F** show the kinetics of total reactive oxygen species (ROS) produced by RAW cells upon challenge with BCG (**C** and **D**) or *M. leprae* (**E** and **F**). *LRRK2* knock out (KO) cells and the nonsynonymous mutations (1628 P/P, 1628 R/P, and 755 L/L) were compared to WT using two-way ANOVA. \**p* < 0.05; ** *p* < 0.01 and *** *p* < 0.001. The experiments were repeated at least three times with similar results. Compared to WT, *LRRK2* KO cells consistently produced less ROS upon bacterial challenge with BCG or *M. leprae* (**C** - **F**). In contrast, there was no significant difference in the kinetics of ROS production following exposure to BCG between WT and LRRK2 755 L/L expressing cells (**D**). Compared to cells expressing LRRK2 WT, cells expressing the LRRK2 1628 P/P variant consistently showed increased ROS production following exposure to both BCG and *M. leprae* (**C, E**). Compared to LRRK2 WT, cells heterozygotic for LRRK2 1628 R/P also displayed higher ROS production upon *M. leprae* stimulation which was not significantly different from the response of homozygous LRRK2 1628 P/P carriers.

Since LRRK2 had been linked to protection from apoptosis, we assessed the impact of the 1628P and 755L mutant LRRK2 proteins on BCG-triggered apoptosis. We infected cells expressing LRRK2 wild type, 1628P/P or 755L/L LRRK2 mutant forms, and LRRK2 knock-out cells with BCG for 24 hours. We then determined in BCG infected and uninfected controls the extent of apoptosis. Following infection with BCG for 24 hrs there was an induction of apoptosis in cells of all lines (Figure 3). Compared to LRRK2 knock-out cells, all other cell lines displayed a reduction in the extent of apoptosis triggered by BCG. Cells expressing wildtype and 755L forms of LRRK2 were indistinguishable by their BCG triggered apoptotic activity. However, cells expressing the 1628P mutant form of LRRK2 showed significant abrogation of apoptosis relative to wildtype cells (Figure 3). Hence, our results revealed a consistent impact of LRRK2 on BCG-induced apoptosis and identified LRRK2 1628P as gain-of-function variant and strong apoptosis reducing factor.

**Fig 3.**
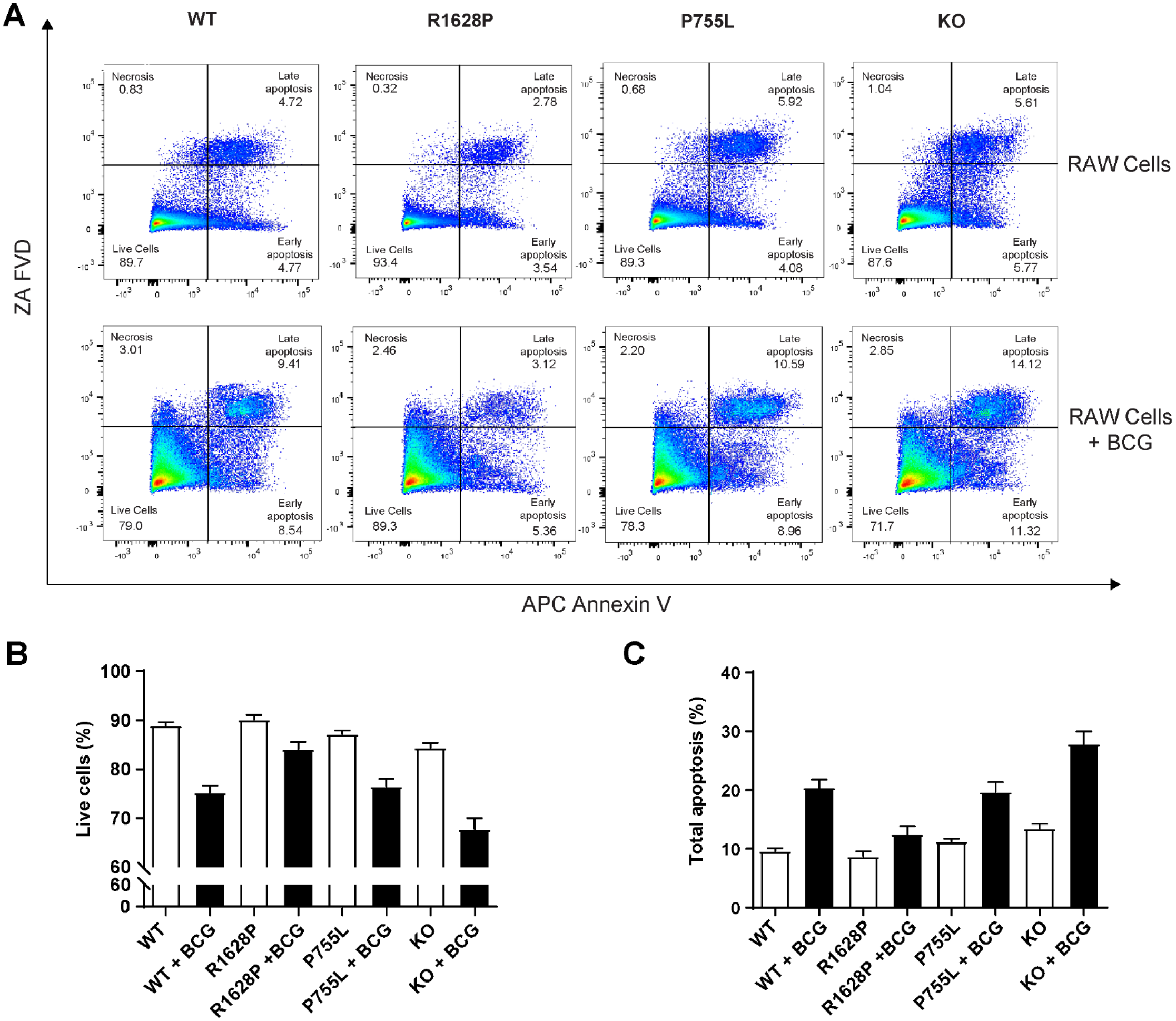
Homozygous LRRK2 1628P/P abrogates apoptosis induced by live BCG. Fluorescence-activated cell sorting (FACS) was used to evaluate the impact of LRRK2 nonsynonymous variants on apoptotic cell death upon BCG challenge. The top of panel A shows the cellular staining as revealed by FACS analyses for RAW cells expression LRRK2 WT and genome-edited proteins. The bottom of panel A shows the same cells after 24 hours co-incubation with live BCG. Panel B denotes the proportion of live cells while panel C indicates the proportion of cells undergoing apoptosis. The bar plots in **A** and **B** display the mean estimate for two experiments done in triplicates with their respective standard errors. There were no significant differences in the proportions of live or apoptotic cells under unstimulated condition (panel **B** and **C**). Infection with BCG triggered a significant (*p* < 0.0001) increase of apoptosis in wildtype, 755L LRRK2 and LRRK2 depleted (KO) cells with a corresponding decline of live cells. In contrast, cells expressing 1628P LRRK2 protein did not respond with a significant increase of dead or apoptotic cells (panel **B** and **C**). When stimulated with BCG, cells depleted of LRRK2 (KO) expressed significantly higher apoptosis when compared to wildtype (*p* = 0.0025), 755L LRRK2 (*p* = 0.0008) or 1628P LRRK2 (*p* < 0.0001) expressing cells (panel **C**).

### Rare nonsynonymous variants in PRKN are T1R-risk factor

For the *PRKN* gene we observed nine missense variants of which two, rs1801474 (S167N) and rs1801582 (V380L), were common and seven were rare including the novel *PRKN* mutation K299R (Figure 1B). *PRKN* rare variants were observed only in T1R-affected cases resulting in strong evidence for gene-wise association of *PRKN* with T1R susceptibility (Figure 1B). To evaluate if the *PRKN* association with T1R was only driven by rare variants we reanalyzed the data removing the two common *PRKN* polymorphisms and observed an approximately one-log drop in strength of association (*p* SKAT-O = 7.4 x10^−5^ to *p* SKAT-O = 1.1 × 10^−3^). Next, we tested if the two common *PRKN* missense variants were associated with T1R in a single-variant model. We observed a trend for association of T1R and the *PRKN* V380L variant (*p* = 0.09; OR (95% CI) = 1.44 (0.94 - 2.20) for 380L carriers), but no association of T1R and S167N (*p* = 0.22; Table S3). To confirm that the V380L variant contributed to the gene-wise association we conditioned the gene-wise model with all nonsynonymous variants of *PRKN* on the V380L mutation. We observed the same drop in the *p* value as in the gene-wise analysis with only rare variants (*p* SKAT-O = 7.4 x10^−5^ to *p*SKAT-O = 5.7 ×10^−3^). This observation indicated that although the rare variants were the main driver of the *PRKN* association with T1R the common V380L variant also contributed to T1R-risk.

The majority of the *PRKN* rare variants found in T1R patients located to or near the Parkin RING domains (Figure 1B). Although the rare variants were dispersed across the linear structure of Parkin, in a 3D model the amino acid substitutions clustered in the central structure of inactive Parkin (Figure 4). Alterations in three Parkin residues (D243N, R275Q and R366W) were predicted to be damaging in all five databases curated (Table S4). Two other substitutions (R42C and K299R) were predicted to impair Parkin function in more than one database while the remaining two rare variants located towards the terminal end of Parkin (R396G and K408R) were predicted to be tolerable (Figure 1B and Table S4). In summary, five Parkin rare variants observed in T1R-affected cases have consistent data supporting a functionally detrimental impact at the protein level.

**Fig 4.**
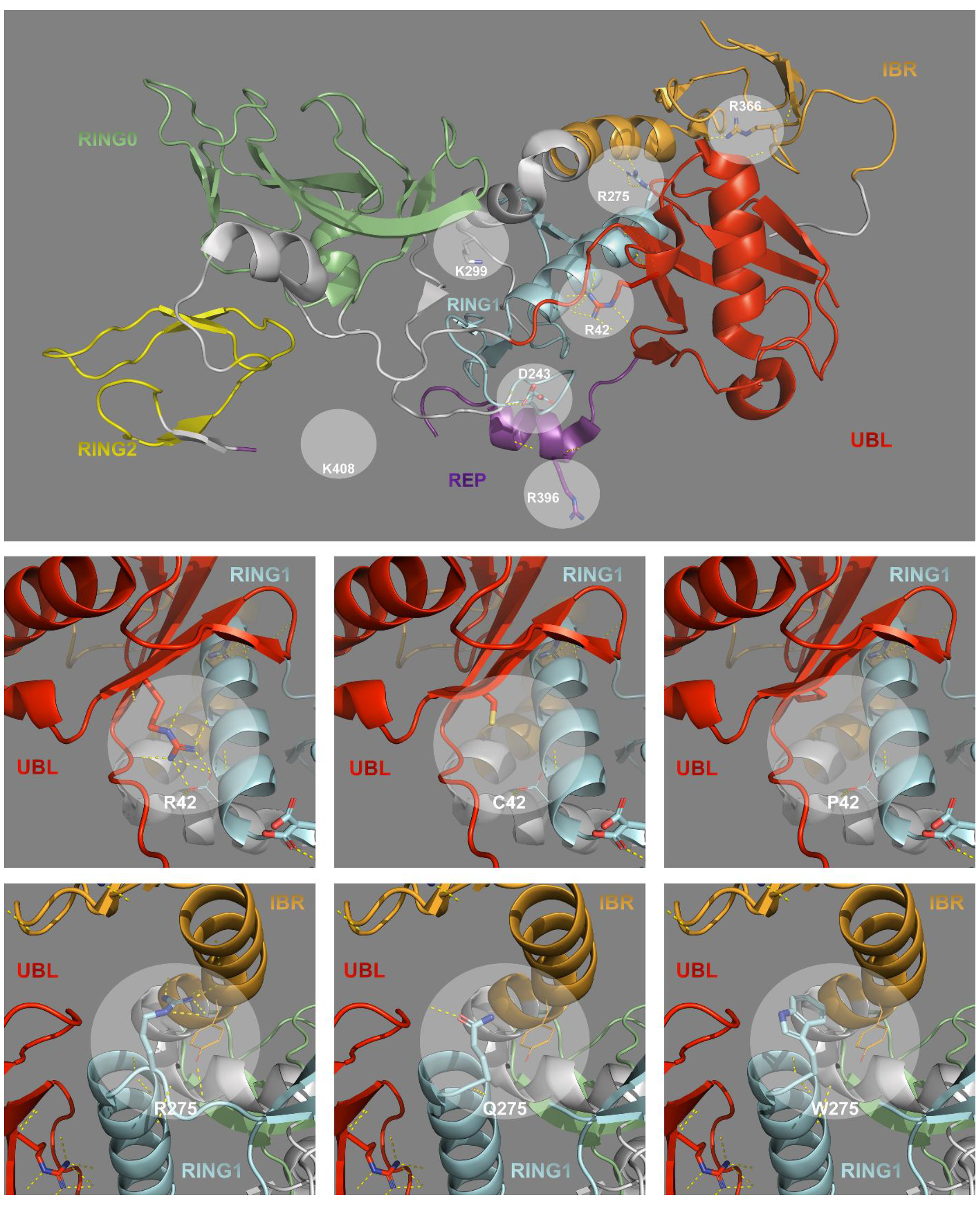
Structural impact of Parkin mutations. At the top, the crystalized structure of Parkin (5C1Z) is given with its respective domains color coded according to Figure 2. The wildtype amino acid residues mutated in T1R-affected subjects are highlighted with circles. Parkin residues 406 to 413 were absent in the crystallography, therefore, the approximate location of the K408R mutation is presented. The central figures represent the wildtype arginine (R) on the left and the variants cysteine (C) at the center and proline (P) to the right for Parkin position 42. At the bottom, the wildtype arginine (R) on the left and the variants glutamine (Q) at the center and tryptophan (W) to the right for Parkin position 275. The diameter of the highlighted circles corresponds to a 5 Armstrong (Å) distance used for the conformation analysis. Interaction between atoms are represented by yellow dotted lines.

### Pleiotropic effect of LRRK2 and PRKN amino acid changes on T1R and Parkinson’s disease

Since mutations in *LRRK2* and *PRKN* are main causes of Parkinson’s disease we assessed if the amino acid substitutions identified in our study were curated in two PD mutation databases ^20,21^. The main evidence for association between *LRRK2* and T1R was provided by variants P755L and R1628P and, while not causally linked with PD, both variants are risk factors for PD (Table 2). Interestingly, antagonistic pleiotropy of the 1628P variant was also observed for PD and Alzheimer’s disease ^22^. For the *PRKN* gene, 5 of the 7 amino acid mutations identified as risk factors for T1R had previously been implicated in PD (Figure 2, Table 2). Moreover, one of the 7 variants was novel and therefore had not been detected in PD or healthy subjects. This shared co-occurrence of amino acid mutations between PD and T1R was strongly non-random (*p* = 8.7 ×10^−4^; Table S5). When we disregarded the novel variant sharing became more significant (1.5 ×10^−4^). Substitutions at Parkin residues 42 and 275 are frequently observed in subjects suffering from PD. The amino acid substitutions R42C observed in oneT1R-affected individual causes the same salt bridge disruption between UBL and RING1 domains as observed for the most common PD R42P mutation (Figure 4). Moreover, the R275Q amino acid substitution affects the interaction between the RING1 and IBR domains of Parkin as observed for the PD R275W mutation (Figure 4). Alterations at the other three Parkin mutations shared between PD and T1R (D243N, R366W and R369G) also result in structural variation (Figure S3).

**Table 2.**
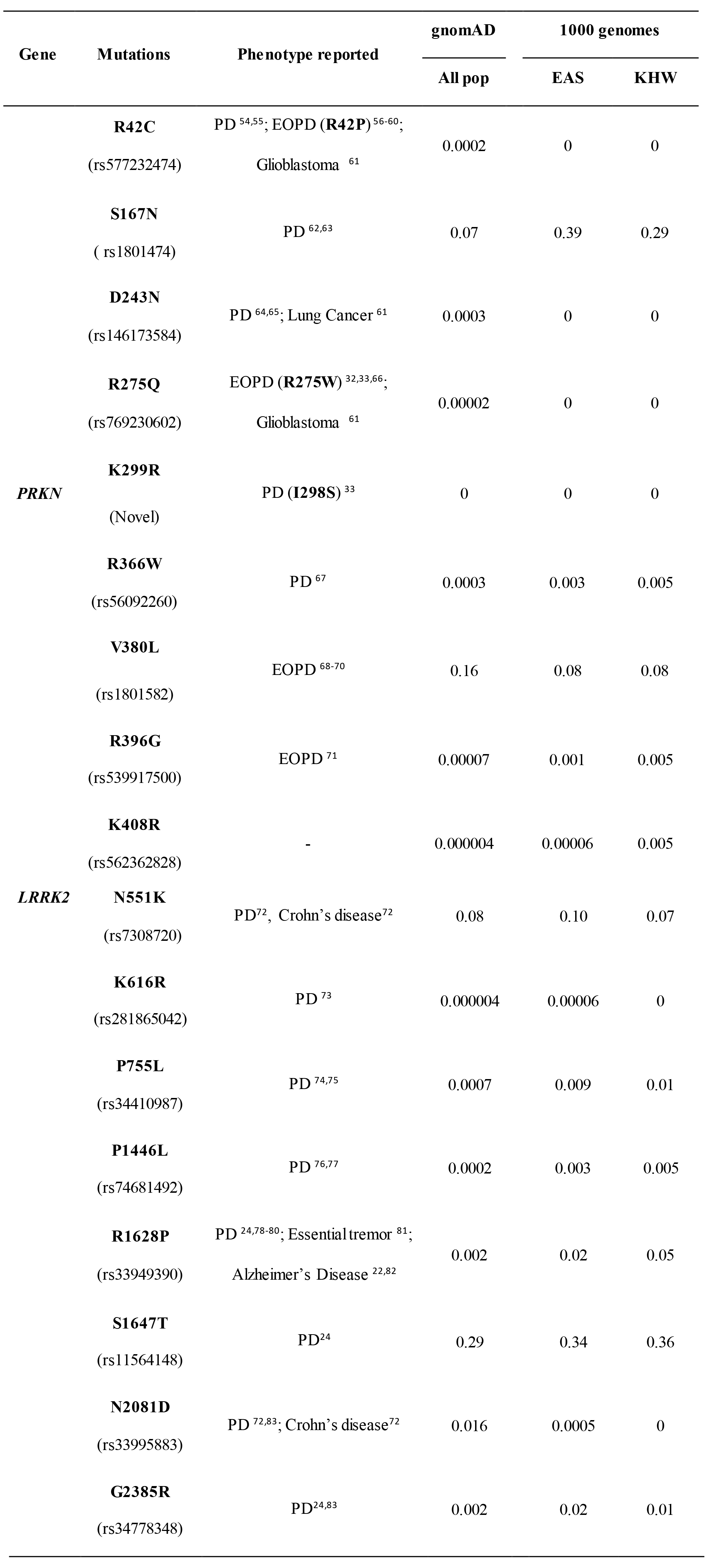
Reported association for variants observed in our study.

## Discussion

Despite the availability of an efficacious treatment for leprosy, the incidence of leprosy has shown only a moderate global decline over the past 15 years ^23^. This unexpected epidemiological situation has given rise to two major areas of research. The first theme concerns the identification of the unknown mode of transmission of the *M. leprae* bacillus. The second theme is focused on the early detection of signs of leprosy and the prevention of nerve damage in leprosy patients ^7^. A main contributor to nerve damage are T1R and the question remains why only a minority of leprosy patients develop T1R and how to identify those who are at risk. While the clinical manifestation of this predisposition requires the trigger by *M. leprae* it is noteworthy that T1R can occur after microbiological cure of leprosy.

Acute episodes of T1R are characterized by a cytokine storm which is indicative of dysregulated inflammatory responses. Prospective transcriptomic analyses of the immune response to *M. leprae* antigen and a genome-wide association study support the central role of host genetics in the break-down of coordinated gene regulation in the T1R phenotype ^12,18^. Additional candidate gene T1R association studies identified T1R susceptibility genes that were investigated in the present manuscript. However, these genetic studies focused on the role of common genetic variants which are more likely to impact on gene expression levels and not on the functional capacity of encoded proteins. Here, we show that for 2 of 7 T1R susceptibility genes, uncommon amino acid mutations that impact protein function are significant contributors to T1R risk that act independently of common regulatory variants. Our findings that a gain-of-function mutation in the LRRK2 protein is protective for T1R while loss of function variants in Parkin are T1R risk factors are consistent with the concept that a genetically controlled mis-communication between pro-and anti-inflammatory host responses underlies the T1R phenotype.

Of the seven T1R candidate genes studied (Table S2), we identified rare and low frequency nonsynonymous variants only in the E3 ligases Parkin and LRRK2 as significantly associated with *M. leprae* triggered pathological inflammation. E3 ligases catalyze the ubiquitination of specific substrates that are involved in a multitude of cellular processes, including protein degradation, transcriptional activity and cellular signaling. The substrate spectrum for the Parkin and LRRK2 is only partly known, and the possible overlap in their substrate landscape remains undefined. However, STRING analysis revealed a high score for interaction of Parkin with LRRK2 suggesting that biological pathways impacted by the two E3 ligases show at least some overlap. As for T1R, our findings validated two key players of nerve damage in leprosy and identified E3 ligases as critical players of this pathology. However, the observed antagonistic pleiotropy of LRRK2 variants in T1R and PD highlights the challenges of host directed therapies that may be aimed at these key mediators of nerve damage in leprosy.

One of the unexpected findings was the observation that the 1628P mutation blocked BCG triggered apoptosis but increased the respiratory burst of RAW cells in response to BCG. These findings are consistent with a previous report that the LRRK2 1628P allele promoted increased kinase activity ^24^. ROS generated during a respiratory burst down-regulate inflammation ^25^, and systemic ROS does induce apoptosis which is generally considered an anti-inflammatory cellular response since apoptotic cells release multiple anti-inflammatory mediators ^26^. However, if apoptotic cells are not cleared efficiently, this may lead to increased inflammation ^27^. This dual role of apoptosis may underlie the antagonistic pleiotropy of 1628P in PD and T1R. It is possible that the reduction of anti—inflammatory molecules resulting from abrogated apoptosis mediated by the 1628 mutant protein is disease-promoting for PD patients while the resulting lower yield of apoptotic debris in leprosy patients may protect against T1R. A previous prospective study of leprosy patients at risk of T1R identified higher pro- and antiinflammatory gene expression in those patients destined to suffer T1R ^12^. This suggested a breakdown in the regulation of the inflammatory response in T1R patients opening the possibility that the 1628P mutation, at least in the periphery, re-establishes the equilibrium between pro-and anti-inflammatory pathways.

Parkin is an evolutionary conserved modulator of innate immunity to infectious agents ^28,29^ and genetic variants of *PRKN* were associated with increased susceptibility to both leprosy and typhoid fever in ethnically diverse populations ^2,30,31^. However, it was not resolved if the role of Parkin in infectious and neurodegenerative disease reflected the promiscuous nature of E3 ligase functions or was based on shared mechanisms of dysfunctional host immunity. Here, we showed that rare amino acid changes associated with T1R in Parkin are predicted loss of function mutations that display a striking overlap with established risk factors for PD. Indeed, five out of the seven Parkin rare variants associated with T1R susceptibility were substitutions at the same amino acid residue as observed in PD while a sixth novel T1R-risk variant (K299R) was adjacent to the annotated I298L and I298S PD mutations (Table 2) ^32,33^. As the shared Parkin mutations are mainly connected to early-onset PD (EOPD) our findings strongly suggested that EOPD shares a mechanistic overlap with T1R, an inflammatory disorder triggered by *M. leprae*.

While the same Parkin mutations are implicated in neurodegeneration in the brain and nerve cell destruction in the periphery, the question remains if related infectious triggers are necessary for the manifestation of both disease phenotypes. Parkin, together with PINK1, is critically important for mitochondrial quality control by modulating mitophagy, a form of autophagy which delivers damaged mitochondria to lysosomes for degradation. However, there is a large body of experimental evidence which extends the role of *PRKN* and *PINK1* in mitochondrial quality control beyond the control of mitophagy ^34^. Given that mitochondria are key players of adaptive and innate immunity, and the suggested role of *Prkn* and *Pink* in autoimmune-mediated neuronal destruction in a mouse model, it is possible that unrecognized infection provides the missing link between impaired mitochondrial quality control and dysregulated neuro-inflammation in EOPD ^35^. The genetic link between EOPD and T1R leads to one additional consideration. Since a majority of T1R cases present within 3 years of diagnosis and treatment of leprosy it seems prudent to consider the role of temporally remote and largely resolved infections as possible triggers for some cases of EOPD. Careful screening for the immunological footprint of such remote infections in EOPD cases may provide additional support for a role of infections in EOPD.

## Online Methods

### Population sample and deep resequencing

A total of 474 leprosy affected subjects were selected from our records of the Vietnamese population. The follow-up for leprosy affected individuals without T1R was greater than 5 years. An Illumina TrueSeq Custom Amplicon v1.5 was used to create libraries containing the exonic sequences of seven targeted genes. Individual indexed libraries were pooled in batches of 96 samples randomly mixing cases and controls for paired-end sequencing using Illumina MiSeq 600 cycles kit v3. Next, the paired-end sequences were aligned to the reference build hg19 of the human genome using a banded Smith-Waterman algorithm from Illumina. In all steps, standard parameters were used except for the exclusion of duplicated read as our approach was based on amplicons. Deep resequencing for the seven genes resulted in a population mean sequence coverage of 262X ranging from 93X for the sample with lowest coverage to 646X for the highest coverage (Figure. S1). Of the 42,265 exonic bases sequenced, 18,354 were in protein coding region. Focusing on coding sequences 92.6% of the bases were covered with more than 30X (Figure. S2).

### Alignment and variant calling

Haplotype caller was used to create individual GVCF files that were combined to perform joint genotyping using the GATK pipeline ^36^. Variants were hard filtered using the suggested GATK thresholds for amplicon sequencing employing the following parameters for SNPs (QD < 2, MQ < 40, SOR > 3, MQRankSum < −12.5 and ReadPosRankSum < −8.0) and for INDELs (QD < 2, SOR > 10, ReadPosRankSum < −20.0 and InbreedingCoeff < −0.8). Additional thresholds were used to ensure the quality of the variant calling by (i) setting as missing genotypes with GQ < 30 and/or with depth of coverage per individual <10X and (ii) excluding variants with a population mean depth of coverage < 30X and call rates < 90%. The function of the variants that passed QC were annotated with ANNOVAR ^37^. Transcribed variants (UTRs, splice sites, synonymous and nonsynonymous variants) and the subset focusing only on codding nonsynonymous variants (missense, stop gain/loss, and frameshifts) were selected for further evaluation.

### Statistical analyses

The enrichment of protein altering variants per gene was evaluated using two different statistical tests: the variable threshold (VT) ^38^ method and the optimal unified burden and SNP-set Kernel Association Test (SKAT-O) ^39^ both implemented in the software EPACTS ^40^. Briefly, VT is a burden test which collapse rare variants in a gene into a single burden variable and test for the cumulative effect of rare variants in the gene by regressing the burden variable on the phenotype. Burden tests are the most powerful rare variant association tests when most variants are causal and the effects are in the same direction. However, in the presence of a mixed of protective and deleterious variants in a gene, the non-burden SKAT is more powerful. To maximize power under both scenarios, SKAT-O adaptively combines a weighted burden and the SKAT statistics. SKAT-O with default parameters was used to weight variants according to their minor allele frequency adding the optional correction for studies with less than 2000 samples. Bonferroni was used for multiple testing correction by considering two tests (SKAT-O and VT) in 7 genes and two conditions (transcribed variants and protein altering) resulting in a cut-off of p < 0.002 to be significant at the gene-wise level. To evaluate if the effect per gene was driven by a particular variant, an univariable analysis was performed for nonsynonymous polymorphisms with MAF > 0.01 under additive model in PLINK ^41^. For the genes associated with T1R in the gene-wise analysis that also presented a variant with *p* < 0.1 in the univariate model a conditional gene-wise association analysis was performed with EPACTS.

A hypergeometric test was used to determine if T1R and PD cases shared substitutions at the same amino-acid residues for the LRRK2 and Parkin proteins more often than expected by chance. As not all amino acids of a protein have the same probability of being mutated in the general population, two databases - (i) dbSNP ^42^ and (ii) gnomAD ^43^ - were accessed to catalog both LRRK2 and Parkin mutated residuals. In the enrichment analysis, only protein mutated residuals with frequency lower than 1% in the curated databases were considered as baseline since common mutations are likely to be present in both T1R and PD samples merely due to their frequency. For instance, in the curated databases Parkin presented 287 out of its 465 amino acids residuals mutated at least once in the general population. The number of Parkin residues associated with PD in the PDmutDB ^20^ and LOVD ^21^ databases was 72. The hypergeometric test calculated the statistical significance of randomly selecting 5 PD-associated residues in the T1R samples when 6 out of the 287 known mutated positions were observed in our population (Table S5). The same test was performed including novel variants for *PRKN* and applied to the *LRRK2* gene.

### Genome-editing with CRISPR/Cas9

For the genome-editing of RAW 264.7 cells with CRISPR/Cas9, the gRNAs for generation of LRRK2 P755L, R1628P and KO cell lines were synthesized by using GeneArt precision gRNA synthesis kit from Thermo Fisher according to the manufacturer’s instruction (Table S6). On the day prior to transfection, RAW264.7 cells were split into a new flask with fresh growth medium such that the cells reach 70-90% confluency the following day. On the day of electroporation, cells were washed with PBS and treated with 0.25% trypsin-EDTA for 8-10 min at 37°C. After neutralization of trypsin, 1 × 10^5^ cells per transfection were transferred to a 1.5 ml microcentrifuge tube. In parallel with the preparation of cells for electroporation, 2μg Cas9 protein and 400 ng gRNA were mixed in 10 μl of resuspension buffer R and incubated at room temperature for 10 min to form stable Cas9-gRNA complex. Prepared cells were resuspended in the buffer R (provided in Neon transfection system 10 μl kit, Thermo Fisher) containing the Cas9-gRNA complex and 50 pmol of donor homology-directed recombination (HDR) templates was added (Table S6). The cell mixture was transferred into a 10 μl Neon tip with Neon pipette and electroporation was performed using the following parameters: pulse voltage 1680 V, pulse width 20 ms and pulse number 1. After electroporation, cells from two Neon tips were transferred to 12-well plates and cultured on growth media for 4 days. Next, the cells were counted and serially diluted to 2 × 10^4^cells/ml, 5 × 10^2^cells/ml and 5 cells/ml. A volume of 200 μl of 5 cells/ml suspension was dispensed to each well of96-well plates. Plates were incubated at 37°C in a 5% CO2. Genomic DNA was isolated from single clone colonies. The donor target region of single clones was Sanger sequenced to check incorporation of the designed variants. Selected RAW264.7 cells were maintained in Dulbecco’s Modified Eagle (DMEM) medium supplemented with 10% fetal bovine serum and 1% streptomycin-penicillin, and incubated in a humidified atmosphere containing 5% CO2 at 37°C. The cells were passaged every 3 days.

### Growth of mycobacteria

BCG Russia cultures were maintained in middlebrook 7H9 medium supplemented with 10% ADC, 0.1% Tween 80, and 0.2% glycerol at 37°C on a roller. On the day of infection to break up large aggregates into single cells, re-suspended BCG was sonicated in a water bath for 20 s × 5 times, followed by passing the BCG through a 22 1/2-G needle 8 times. Bacterial load was determined by plating serial 10-fold dilutions of BCG and CFUs were estimated from colony counts after at least 3 weeks of incubation. The Thai-53 isolate *M. leprae* was harvested from both hind foot pads of athymic nude mice (Envigo, USA) inoculated 5-7 months previously with 3 × 10^7^ bacilli. The harvested bacilli were enumerated by direct count according to Shepard’s method and held overnight at 4 – 8 °C pending quality control testing for contamination and viability. Viability of the *M. leprae* suspension was measured by determining the rate of 14C-palmitic acid oxidation to 14CO2 by radiorespirometry as described previously ^44^. A 24hr count of at least 6000cpm was considered viable. Freshly harvested bacilli were always employed in experiments within 24-48hrs of harvest.

### Western Blot analysis

Total cellular lysates were resolved on a 4 to 12% Tris-Glycine gel (Bio-Rad) and electrophoretically transferred to polyvinylidene difluoride membranes (Bio-Rad). The membranes were blocked with 5% BSA in TBS-T (Tris-buffered saline-0.1% Tween 20) for 1 h at room temperature, a procedure followed by incubation with primary antibodies overnight at 4°C. A rabbit monoclonal antibody against LRRK2 (Abcam) was used at 1:1,000 dilution. A mouse anti-GAPDH monoclonal antibody (Thermo Fisher) was used at a 1:10,000 dilution. Upon extensive washing, the membrane was developed with enhanced chemiluminescence detection reagents (Bio-Rad), followed by imaging using a ChemiDoc Touch imaging system (Bio-Rad). Experiments were performed in triplicates.

### Reactive oxygen species (ROS) detection

Different cell lines were seeded in 96-well plates at a concentration of 3×10^4^ cells per well and stimulated with IFN-γ (100ng/ml) for 24 hours. Cells were then infected with BCG-Russia or*M. leprae* at an MOI of 10:1. At indicated time points following infection, ROS production was detected using ROS-ID total ROS detection kit (Enzo life science) according to the manufacturer’s instructions. Briefly, cells were carefully washed with 200 μl/well of 1X wash buffer. Following wash buffer removal, 100 μl/well of ROS detection mix (4 μl of 5mM oxidative stress detection reagent /10 ml of 1X wash buffer) was added prior to incubation of plates in a humidified incubator (37°C, 5% CO_2_) for 30 min. Readings were acquired at wavelengths 488/520nm on a plate reader. Each experiment was performed at least three times, each in triplicate.

### FACS analysis of apoptosis

One the day prior to infection, RAW264.7 cells expressing LRRK2 wildtype or mutant protein or depleted for *LRRK2* gene expression by knock-out were separately cultured at a concentration of 6 x10^5^/well in a 6-well plate for 16-18 hrs. The cells were then infected with or without BCG (MOI 10:1) for 24 hours followed by apoptosis analysis using Annexin V-APC and Zombie Aqua TM fixable viability dye as detailed in the Biolegend manual. Briefly, cells were washed twice with cold PBS and incubated with 1ml of 5 mM EDTA/PBS on ice for 30 minutes followed by pipetting to detach and dissociate the cells. Finally, cells were resuspended in 200μL of 1X Binding Buffer and immediately collected for flow cytometry with BD FACSCanto II (BD Biosciences). The data were analyzed on FlowJo^®^ v10.4.2 (FlowJo, LLC) with viability and Annexin V single stains as FMOs.

### Database curation

The predicted impact at the protein level for nonsynonymous variants described in our study was assessed in five databases as in previous studies ^45,46^. The algorithms used for the predictions were PolyPhen2 HumDiv and HumVar ^47^, SIFT ^48^, LRT ^49^, Mutation Taster ^50^ and CADD ^51^. To evaluate if rare variants associated with T1R were also reported in PD cases, we used the PDmutDB (http://www.molgen.vib-ua.be/PDMutDB/) ^20^ and the Leiden Open Variation Database (LOVD) for Parkinson’s Disease (http://grenada.lumc.nl/LOVD2/TPI/home.php) ^21^. To evaluate the frequency of the mutations of the targeted genes we used the dbSNP v147 (https://www.ncbi.nlm.nih.gov/snp) ^42^ and gnomAD (http://gnomad.broadinstitute.org/) ^43^ databases.

### Spatial modeling of Parkin mutations

To visualize the spatial localization of mutations we used a crystallography model of the inactive Parkin (PDB:5C1Z) ^52^. The structural impact of amino acid substitutions in Parkin was predicted with PyMol ^53^. Briefly, rearrangements in Parkin conformation were estimated for amino acids in a five Armstrong (Å) radius from the most likely rotamer of the mutated residuals. The impact in polarity and hydrogen bounding was also assessed for amino acids surrounding the mutated site. One amino acid was mutated at a time with the reference allele observed in the crystallography kept for all other positions. The impact of mutations in Parkin residues 357 to 360, 383 to 390 and 406 to 413 could not be assessed, including the mutation at position 408 identified in our study.

## Acknowledgments

The authors thank all leprosy patients who participated in this study and members of the Schurr lab for many helpful discussions. We thank Kai Sheng for technical support. This work was supported by a Foundation grant from the Canadian Institutes of Health Research (CIHR) to ES (FDN-143332). The provision of *M. leprae* was supported by the NIAID Interagency Agreement IAA 15006-004. This research was supported through resource allocation in the Guillimin high performance computing cluster by Compute Canada (www.computecanada.ca and Calcul Québec (www.calculquebec.ca).

## Author contributions

Conceptualization: VMF, AC, AA, LA and ES; Data curation: VMF, AC and MO; Formal analyses: VMF, YZX, GL and ES; Funding acquisition: ES; Wet lab experiments: VMF, YZX, GC, ST and MO; Resources: NVT, MO, VHT, RL and LA; Writing – original draft: VMF, YZX and ES; Writing – review and editing: GL, AC, AA, LA, RL, LA and MO.

## Competing interests

The authors declare no conflict of interest.

## Supplementary Materials

**Fig S1.**
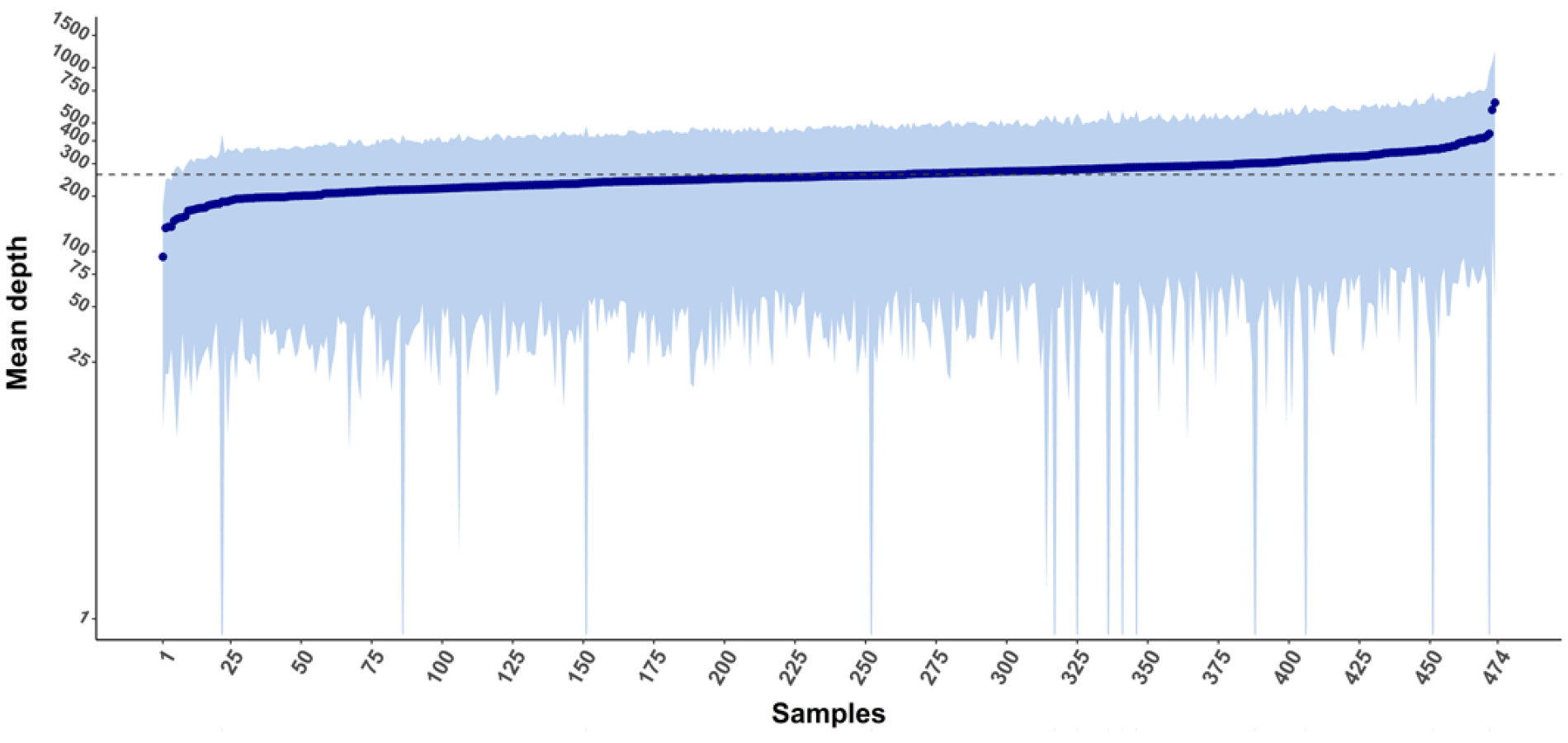
Mean depth of coverage per sample in the T1R population. The depth of coverage (average number of reads per sequenced base) was plotted as a log scale in the y-axis for the 474 studied subjects in the x-axis. Dark blue circles represent the depth of coverage per individual and the light blue shade indicates the standard deviation of the mean. The horizontal dotted line marks the population mean coverage of 262.5X.

**Fig S2.**
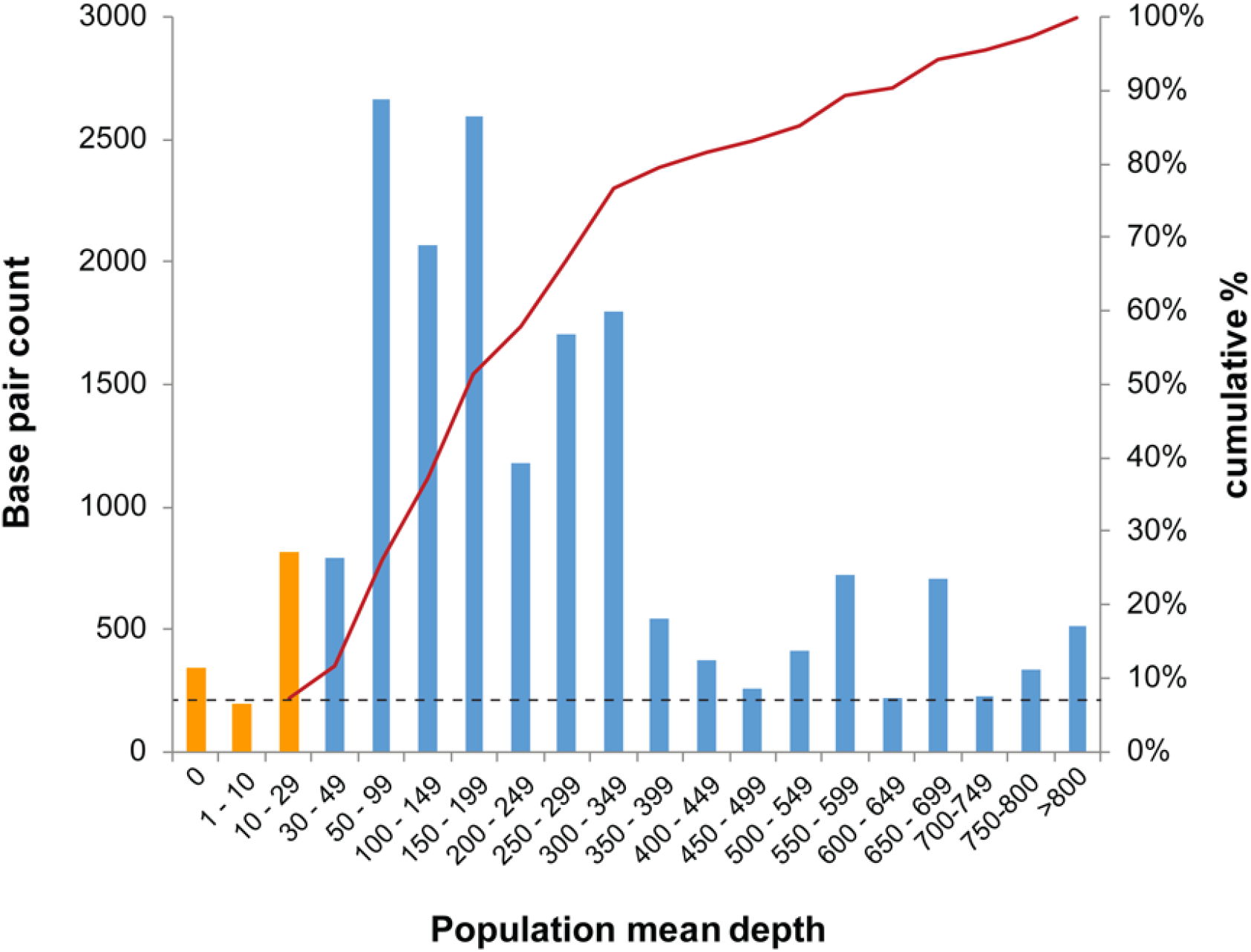
Population mean depth of coverage for translated bases in T1R targeted genes. The total number of coding bases of T1R targeted genes is plotted on the left y-axis according to bins of population mean depth of coverage on the x-axis. The orange bars represent the number of bases that had less than 30X reads while blue bars were bases covered above 30X. The cumulative proportion of base coverage is plotted on the right y-axis. A dotted horizontal line represents the cut off of 30X while the red line indicates the cumulative proportion of bases above 30X.

**Fig S3.**
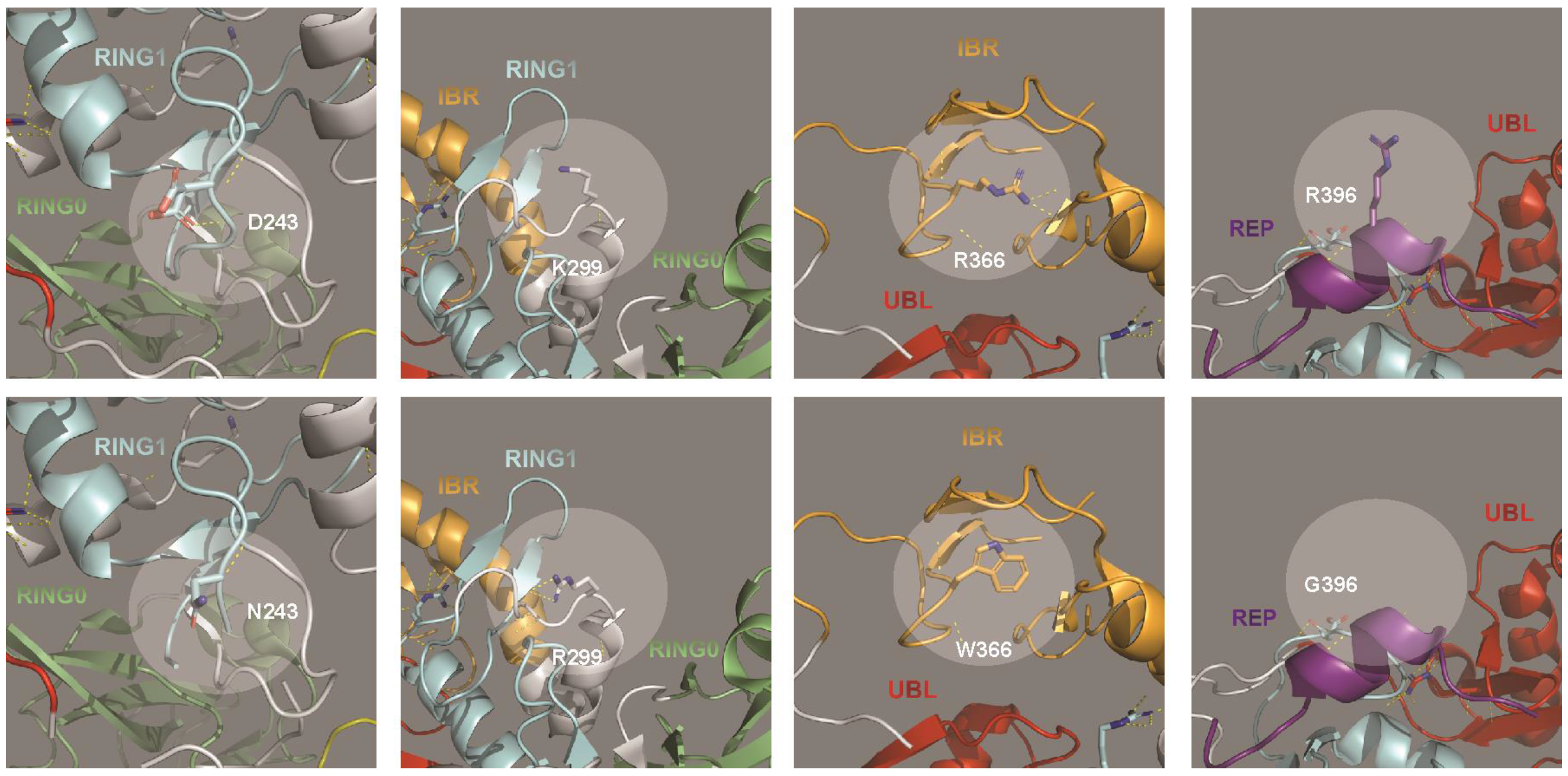
Structural impact of additional Parkin mutations. The four images on the left represent the wildtype alleles for Parkin at positions 243, 299, 366 and 396 while the right images represent the amino acid substitutions observed in T1R-affected cases. The diameter of the highlighted circles corresponds to a 5 Armstrong (Å) distance used for the conformation analysis. Interaction between atoms are represented by yellow dotted lines.

**Table S1.** Epidemiological data of the studied sample.

|  | **T1R affected** | **T1R free** |
| --- | --- | --- |
|  | 237 | 237 |
| **Age at leprosy diagnosis** |  |  |
| Mean (S.D.) | 19.8 (5.4) | 20.5 (5.1) |
| **Gender** |  |  |
| Male | 165 | 165 |
| Female | 72 | 72 |
| **Clinical Subtype** |  |  |
| TT | - | - |
| BT | 50 | 49 |
| BB | 102 | 102 |
| BL | 85 | 86 |
| LL | - | - |

T1R, type-1 reaction; SD, standard deviation; TT, tuberculoid-tuberculoid; BT, borderline-tuberculoid; BB, borderline-borderline; BL, borderline-lepromatous; LL, lepromatous-lepromatous. The follow-up for leprosy affected individuals without T1R was greater than 5 years.

**Table S2.** Exonic variants in the T1R targeted genes.

|  | *TLR1* | *TLR2* | *NOD2* | *TNFSF15* | *TNFSF8* | *LRRK2* | *PRKN* | *Total* |
| --- | --- | --- | --- | --- | --- | --- | --- | --- |
| UTR5' | 6 | 1 | 0 | 1 | 1 | 0 | 0 | 9 |
| Synonymous | 4 | 6 | 7 | 2 | 2 | 13 | 1 | 35 |
| Nonsynonymous |  |  |  |  |  |  |  |  |
| Missense | 13 | 7 | 8 | 2 | 0 | 19 | 9 | 58 |
| Frameshift deletion | 0 | 0 | 1 | 0 | 0 | 0 | 0 | 1 |
| Frameshift insertion | 0 | 0 | 1 | 0 | 0 | 1 | 0 | 2 |
| Nonframeshift deletion | 0 | 1 | 0 | 0 | 0 | 0 | 0 | 1 |
| Stop gain | 0 | 1 | 0 | 0 | 0 | 0 | 0 | 1 |
| UTR3' | 0 | 3 | 8 | 19 | 34 | 15 | 13 | 92 |
| **Total** | 23 | 19 | 25 | 24 | 37 | 48 | 23 | 199 |

**Table S3.**
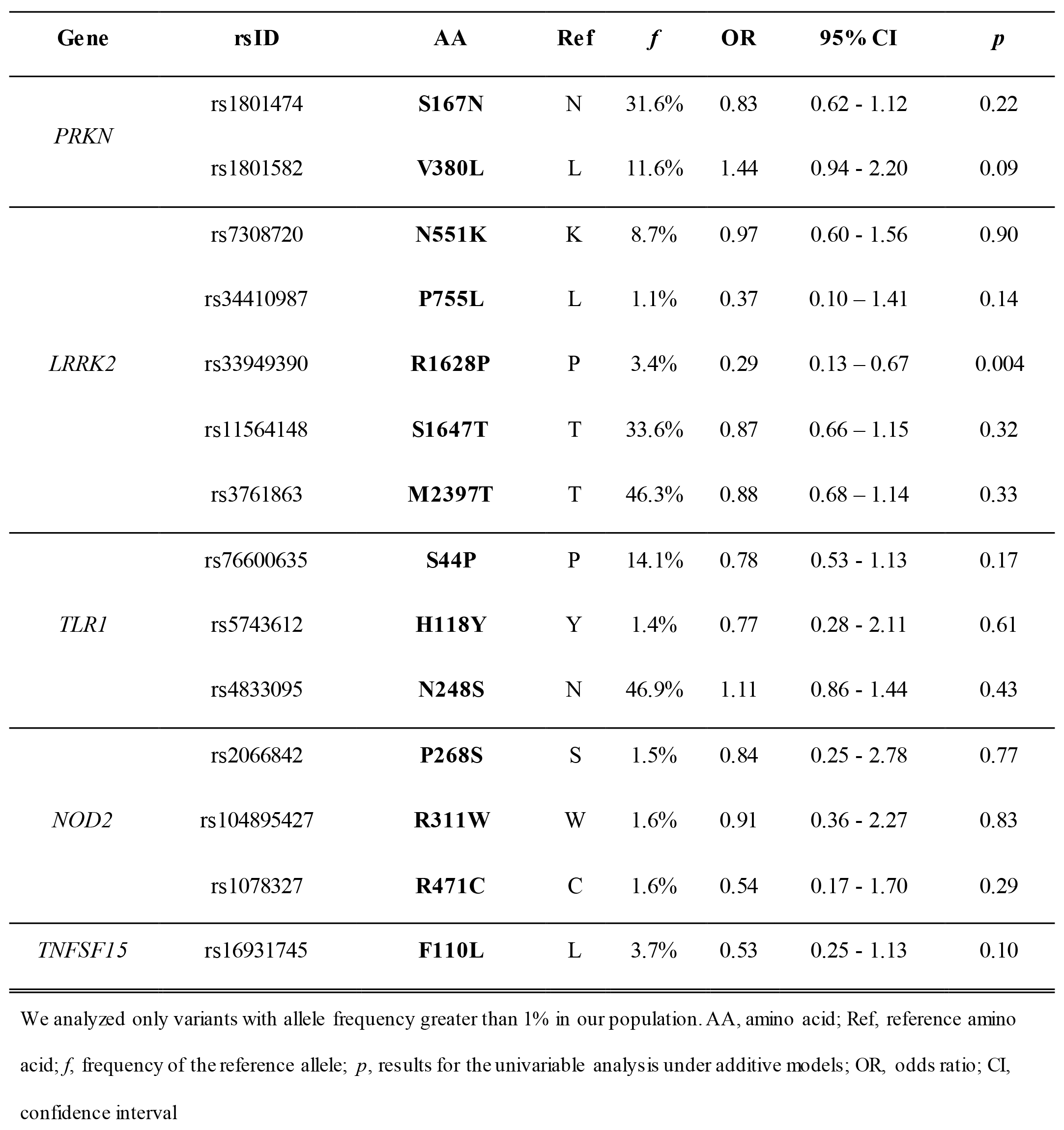
Univariable analysis of amino-acid changes in T1R risk genes.

| Gene | rsID | AA | Ref | *f* | OR | 95% CI | *p* |
| --- | --- | --- | --- | --- | --- | --- | --- |
| *PRKN* | rs1801474 | **S167N** | N | 31.6% | 0.83 | 0.62 - 1.12 | 0.22 |
|  | rs1801582 | **V380L** | L | 11.6% | 1.44 | 0.94 - 2.20 | 0.09 |
| *LRRK2* | rs7308720 | **N551K** | K | 8.7% | 0.97 | 0.60 - 1.56 | 0.90 |
|  | rs34410987 | **P755L** | L | 1.1% | 0.37 | 0.10 – 1.41 | 0.14 |
|  | rs33949390 | **R1628P** | P | 3.4% | 0.29 | 0.13 – 0.67 | 0.004 |
|  | rs11564148 | **S1647T** | T | 33.6% | 0.87 | 0.66 – 1.15 | 0.32 |
|  | rs3761863 | **M2397T** | T | 46.3% | 0.88 | 0.68 – 1.14 | 0.33 |
| *TLR1* | rs76600635 | **S44P** | P | 14.1% | 0.78 | 0.53 - 1.13 | 0.17 |
|  | rs5743612 | **H118Y** | Y | 1.4% | 0.77 | 0.28 - 2.11 | 0.61 |
|  | rs4833095 | **N248S** | N | 46.9% | 1.11 | 0.86 - 1.44 | 0.43 |
| *NOD2* | rs2066842 | **P268S** | S | 1.5% | 0.84 | 0.25 - 2.78 | 0.77 |
|  | rs104895427 | **R311W** | W | 1.6% | 0.91 | 0.36 - 2.27 | 0.83 |
|  | rs1078327 | **R471C** | C | 1.6% | 0.54 | 0.17 - 1.70 | 0.29 |
| *TNFSF15* | rs16931745 | **F110L** | L | 3.7% | 0.53 | 0.25 - 1.13 | 0.10 |

We analyzed only variants with allele frequency greater than 1% in our population. AA, amino acid; Ref, reference amino acid; *f*, frequency of the reference allele; *p*, results for the univariable analysis under additive models; OR, odds ratio; CI, confidence interval

**Table S4.** Annotation of protein altering variants identified in the T1R targeted genes. Large file that could not be embedded in the manuscript and was submitted separately.

**Table S5.** Parkinson’s disease and T1R affected subjects share amino acid substitution at the same Parkin residues.

| *PRKN* | T1R<br>variant | No T1R<br>variant | *p* |
| --- | --- | --- | --- |
| PD variant | 5 | 67 | 1.5 x 10<sup>-4</sup> |
| No PD variant | 1 | 214 |  |
| Total | 6* | 287 |  |

Parkin mutated residues observed in Parkinson's disease cases were extracted from the PDmutDB and LOVD database. Mutations in Parkin residues were accounted only once even if different amino acid substitutions were cataloged for a given position. Substitutions in Parkin residues 167 and 380 were not included in the analyses due to their high frequency in the general population. A hypergeometric test was used to investigate the pleiotropic enrichment. \*If the novel variant observed in a T1R case by our study was to be included as "No PD" variant the pleiotropic effect would still remain highly significant ( $p = 8.7 \times 10^{-4}$ ).

**Table S6.**
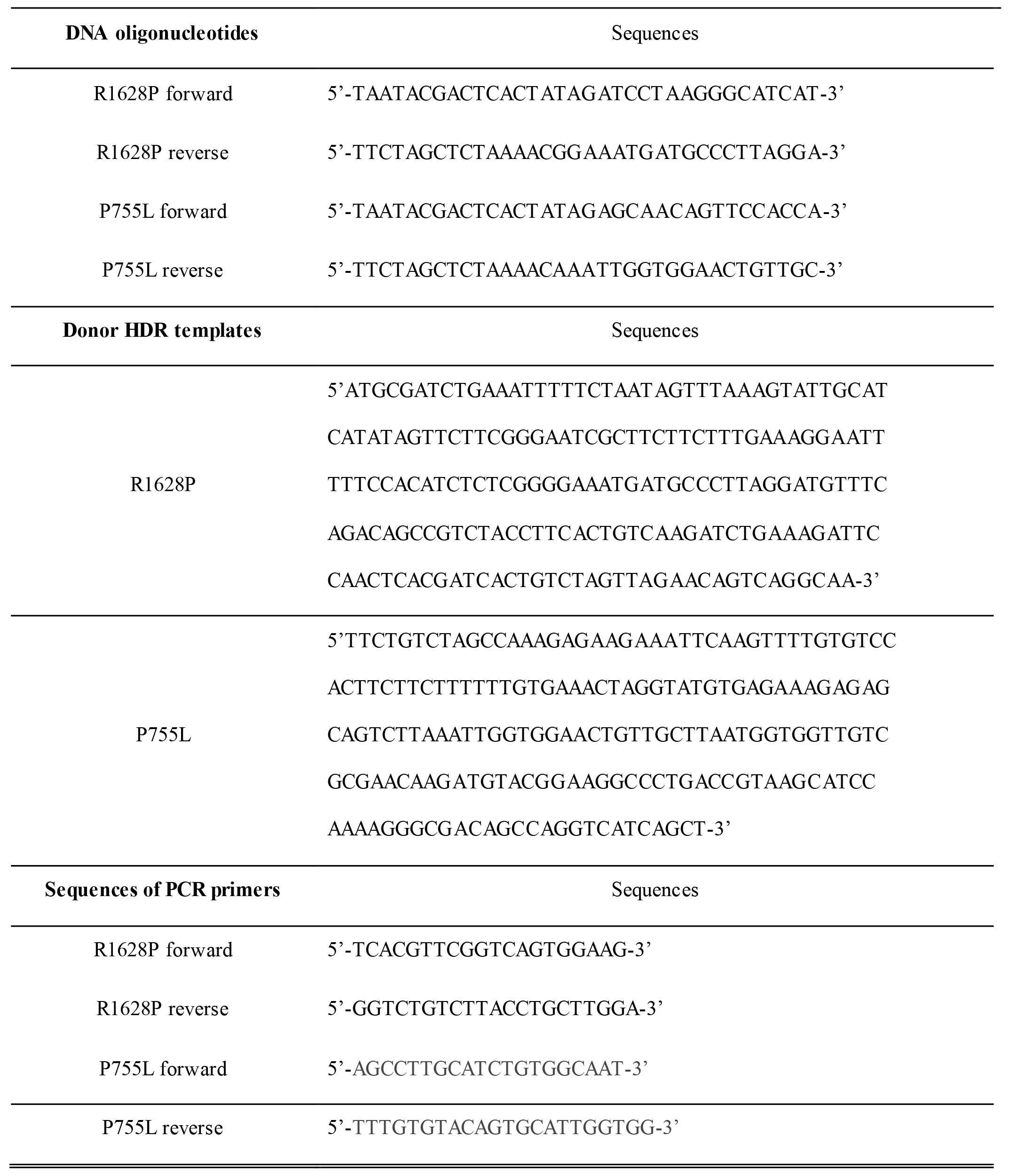
Oligonucleotides used in the CRISPR/Cas9 constructs.

## References

1. Fava V.M. & Schurr E. The complexity of the host genetic contribution to the human response to Mycobacterium leprae. in The International Textbook of Leprosy (eds. Scollard, D.M. & Gillis, T.P.) (American Leprosy Mission, http://www.internationaltextbookofleprosy.org/, 2016).

2. Mira, M.T., et al. Susceptibility to leprosy is associated with PARK2 and PACRG. Nature 427, 636–640 (2004).

3. Alcais, A., et al. Stepwise replication identifies a low-producing lymphotoxin-[alpha] allele as a major risk factor for early-onset leprosy. Nat Genet 39, 517–522 (2007).

4. Zhang, F.R., et al. Genomewide association study of leprosy. N Engl J Med 361, 2609–2618 (2009).

5. Zhang, F., et al. Identification of two new loci at IL23R and RAB32 that influence susceptibility to leprosy. Nat Genet 43, 1247–1251 (2011).

6. Wang, Z., et al. A large-scale genome-wide association and meta-analysis identified four novel susceptibility loci for leprosy. Nature communications 7, 13760 (2016).

7. WHO. Global leprosy strategy 2016-2020: accelerating towards a leprosy-free world - 2016 operational manual. in Global leprosy strategy 2016—2020: accelerating towards a leprosy-free world— 2016 operational manual. (ed. Cooreman, E.A.) 62 (WHO, 2016).

8. Fava, V., et al. Genetics of leprosy reactions: an overview. Mem Inst Oswaldo Cruz 107 Suppl 1, 132–142 (2012).

9. Kahawita, I.P., Walker, S.L. & Lockwood, D.N.J. Leprosy type 1 reactions and erythema nodosum leprosum. Anais Brasileiros de Dermatologia 83, 75–82 (2008).

10. Ranque, B., et al. Age is an important risk factor for onset and sequelae of reversal reactions in Vietnamese patients with leprosy. Clin Infect Dis 44, 33–40 (2007).

11. Sousa, A.L., et al. Genetic and immunological evidence implicates interleukin 6 as a susceptibility gene for leprosy type 2 reaction. J Infect Dis 205, 1417–1424 (2012).

12. Orlova, M., et al. Gene set signature of reversal reaction type I in leprosy patients. PLoS Genet 9, e1003624 (2013).

13. Misch, E.A., et al. Human TLR1 deficiency is associated with impaired mycobacterial signaling and protection from leprosy reversal reaction. PLoS Negl Trop Dis 2, e231 (2008).

14. Bochud, P.Y., et al. Toll-like receptor 2 (TLR2) polymorphisms are associated with reversal reaction in leprosy. J Infect Dis 197, 253–261 (2008).

15. Berrington, W.R., et al. Common polymorphisms in the NOD2 gene region are associated with leprosy and its reactive states. J Infect Dis 201, 1422–1435 (2010).

16. Fava, V.M., Sales-Marques, C., Alcais, A., Moraes, M.O. & Schurr, E. Age-Dependent Association of TNFSF15/TNFSF8 Variants and Leprosy Type 1 Reaction. Front Immunol 8, 155 (2017).

17. Fava, V.M., et al. Association of TNFSF8 regulatory variants with excessive inflammatory responses but not leprosy per se. J Infect Dis 211, 968–977 (2015).

18. Fava, V.M., et al. A genome wide association study identifies a lncRna as risk factor for pathological inflammatory responses in leprosy. PLoS Genet 13, e1006637 (2017).

19. Fava, V.M., et al. A Missense LRRK2 Variant Is a Risk Factor for Excessive Inflammatory Responses in Leprosy. PLoS Negl Trop Dis 10, e0004412 (2016).

20. Cruts, M., Theuns, J. & Van Broeckhoven, C. Locus-specific mutation databases for neurodegenerative brain diseases. Human mutation 33, 1340–1344 (2012).

21. Fokkema, I.F., et al. LOVD v.2.0: the next generation in gene variant databases. Human mutation 32, 557–563 (2011).

22. Li, H.L., et al. The LRRK2 R1628P variant plays a protective role in Han Chinese population with Alzheimer’s disease. CNS Neurosci Ther 19, 207–215 (2013).

23. WHO. Global leprosy update, 2017: reducing the disease burden due to leprosy Wkly Epidemiol Rec 93, 445–456 (2018).

24. Tan, E.K., et al. Multiple LRRK2 variants modulate risk of Parkinson disease: a Chinese multicenter study. Human mutation 31, 561–568 (2010).

25. Luo, B., et al. Phagocyte respiratory burst activates macrophage erythropoietin signalling to promote acute inflammation resolution. Nature communications 7, 12177 (2016).

26. Zhang, L., et al. Redox signaling: Potential arbitrator of autophagy and apoptosis in therapeutic response. Free Radic Biol Med 89, 452–465 (2015).

27. Yang, Y., Jiang, G., Zhang, P. & Fan, J. Programmed cell death and its role in inflammation. Mil Med Res 2, 12 (2015).

28. Manzanillo, P.S., et al. The ubiquitin ligase parkin mediates resistance to intracellular pathogens. Nature 501, 512–516 (2013).

29. Behr M.A. & Schurr E. Cell biology: A table for two. Nature 501, 498–499 (2013).

30. Ali, S., et al. PARK2/PACRG polymorphisms and susceptibility to typhoid and paratyphoid fever. Clin Exp Immunol 144, 425–431 (2006).

31. Chopra, R., et al. Mapping of PARK2 and PACRG overlapping regulatory region reveals LD structure and functional variants in association with leprosy in unrelated indian population groups. PLoS Genet 9, e1003578 (2013).

32. Wang, Y., et al. Risk of Parkinson disease in carriers of parkin mutations: estimation using the kin-cohort method. Arch Neurol 65, 467–474 (2008).

33. Lesage, S., et al. Rare heterozygous parkin variants in French early-onset Parkinson disease patients and controls. J Med Genet 45, 43–46 (2008).

34. Mouton-Liger, F., Jacoupy, M., Corvol, J.C. & Corti, O. PINK1/Parkin-Dependent Mitochondrial Surveillance: From Pleiotropy to Parkinson’s Disease. Front Mol Neurosci 10, 120 (2017).

35. Matheoud, D., et al. Parkinson’s Disease-Related Proteins PINK1 and Parkin Repress Mitochondrial Antigen Presentation. Cell 166, 314–327 (2016).

36. McKenna, A., et al. The Gsnome Analysis Toolkit: a MapReduce framework for analyzing next-generation DNA sequencing data. Genome Res 20, 1297–1303 (2010).

37. Wang, K., Li, M. & Hakonarson, H. ANNOVAR: functional annotation of genetic variants from high-throughput sequencing data. Nucleic Acids Res 38, e164 (2010).

38. Price, A.L., et al. Pooled association tests for rare variants in exon-resequencing studies. Am J Hum Genet 86, 832–838 (2010).

39. Lee, S., et al. Optimal unified approach for rare-variant association testing with application to small-sample case-control whole-exome sequencing studies. Am J Hum Genet 91, 224–237 (2012).

40. Kang H.M. & Abecasis G. EPACTS: Efficient and Parallelizable Association Container Toolbox (2017).

41. Purcell, S., et al. PLINK: a tool set for whole-genome association and population-based linkage analyses. Am J Hum Genet 81, 559–575 (2007).

42. Sherry, S.T., et al. dbSNP: the NCBI database of genetic variation. Nucleic Acids Res 29, 308–311 (2001).

43. Karczewski, K.J., et al. The ExAC browser: displaying reference data information from over 60 000 exomes. Nucleic Acids Res 45, D840–D845 (2017).

44. Lahiri R. & Adams L.B. Cultivation and viability determination of Mycobacterium leprae. in The International Textbook of Leprosy (eds. Scollard, D.M. & Gillis, T.P.) (American Leprosy Mission, http://www.internationaltextbookofleprosy.org/, 2016).

45. Marouli, E., et al. Rare and low-frequency coding variants alter human adult height. Nature 542, 186–190 (2017).

46. Purcell, S.M., et al. A polygenic burden of rare disruptive mutations in schizophrenia. Nature 506, 185–190 (2014).

47. Adzhubei, I.A., et al. A method and server for predicting damaging missense mutations. Nature methods 7, 248–249 (2010).

48. Vaser R., Adusumalli S., Leng S.N., Sikic M. & Ng, P.C. SIFT missense predictions for genomes. Nature protocols 11, 1–9 (2016).

49. Liu, X., Jian, X. & Boerwinkle, E. dbNSFP: a lightweight database of human nonsynonymous SNPs and their functional predictions. Human mutation 32, 894–899 (2011).

50. Schwarz, J.M., Cooper, D.N., Schuelke, M. & Seelow, D. MutationTaster2: mutation prediction for the deep-sequencing age. Nature methods 11, 361–362 (2014).

51. Kircher, M., et al. A general framework for estimating the relative pathogenicity of human genetic variants. Nat Genet 46, 310–315 (2014).

52. Kumar, A., et al. Disruption of the autoinhibited state primes the E3 ligase parkin for activation and catalysis. EMBO J 34, 2506–2521 (2015).

53. Schrodinger LLC. The PyMOL Molecular Graphics System, Version 1.8. (2015).

54. Chaudhary, S., et al. Parkin mutations in familial and sporadic Parkinson’s disease among Indians. Parkinsonism Relat Disord 12, 239–245 (2006).

55. Biswas, A., et al. Molecular pathogenesis of Parkinson’s disease: identification of mutations in the Parkin gene in Indian patients. ParkinsonismRelatDisord 12, 420–426 (2006).

56. Terreni, L., Calabrese, E., Calella, A.M., Forloni, G. & Mariani, C. New mutation (R42P) of the parkin gene in the ubiquitinlike domain associated with parkinsonism. Neurology 56, 463–466 (2001).

57. Bertoli-Avella, A.M., et al. Novel parkin mutations detected in patients with early-onset Parkinson’s disease. Mov Disord 20, 424–431 (2005).

58. Clark, L.N., et al. Case-control study of the parkin gene in early-onset Parkinson disease. Arch Neurol 63, 548–552 (2006).

59. Pellecchia, M.T., et al. Parkinsonism and essential tremor in a family with pseudo-dominant inheritance of PARK2: an FP-CIT SPECT study. Mov Disord 22, 559–563 (2007).

60. Macedo, M.G., et al. Genotypic and phenotypic characteristics of Dutch patients with early onset Parkinson’s disease. Mov Disord 24, 196–203 (2009).

61. Veeriah, S., et al. Somatic mutations of the Parkinson’s disease-associated gene PARK2 in glioblastoma and other human malignancies. Nat Genet 42, 77–82 (2010).

62. Mellick, G.D., et al. The parkin gene S/N167 polymorphism in Australian Parkinson’s disease patients and controls. Parkinsonism Relat Disord 7, 89–91 (2001).

63. Satoh J. & Kuroda Y. Association of codon 167 Ser/Asn heterozygosity in the parkin gene with sporadic Parkinson’s disease. Neuroreport 10, 2735–2739 (1999).

64. Romito, L.M., et al. High frequency stimulation of the subthalamic nucleus is efficacious in Parkin disease. J Neurol 252, 208–211 (2005).

65. Lohmann, E., et al. Genetic bases and phenotypes of autosomal recessive Parkinson disease in a Turkish population. Eur J Neurol 19, 769–775 (2012).

66. Sironi, F., et al. Parkin analysis in early onset Parkinson’s disease. Parkinsonism Relat Disord 14, 326–333 (2008).

67. Wang, M., et al. Polymorphism in the parkin gene in sporadic Parkinson’s disease. Ann Neurol 45, 655–658 (1999).

68. Lucking, C.B., et al. Coding polymorphisms in the parkin gene and susceptibility to Parkinson disease. Arch Neurol 60, 1253–1256 (2003).

69. Gaweda-Walerych, K., et al. PARK2 variability in Polish Parkinson’s disease patients–interaction with mitochondrial haplogroups. Parkinsonism Relat Disord 18, 520–524 (2012).

70. Zhang, Y., Wang, Z.Z. & Sun, H.M. A meta-analysis of the relationship of the Parkin p.Val380Leu polymorphism to Parkinson’s disease. American journal of medical genetics. Part B, Neuropsychiatric genetics: the official publication of the International Society of Psychiatric Genetics 162B, 235–244 (2013).

71. Wu, R.M., et al. Parkin mutations and early-onset parkinsonism in a Taiwanese cohort. Arch Neurol 62, 82–87 (2005).

72. Hui, K.Y., et al. Functional variants in the LRRK2 gene confer shared effects on risk for Crohn’s disease and Parkinson’s disease. Sci Transl Med 10(2018).

73. Wang, L., et al. A novel LRRK2 mutation in a mainland Chinese patient with familial Parkinson’s disease. Neurosci Lett 468, 198–201 (2010).

74. Tomiyama, H., et al. LRRK2 P755L variant in sporadic Parkinson’s disease. Journal of human genetics 53, 1012–1015 (2008).

75. Wu, T., et al. A novel P755L mutation in LRRK2 gene associated with Parkinson’s disease. Neuroreport 17, 1859–1862 (2006).

76. Tomiyama, H., et al. Mutation analysis of LRRK2 exon 31 and a novel P1446L mutation in Asian familial Parkinson’s disease. Movement Disord 21, S575–S575 (2006).

77. Zabetian, C.P., et al. LRRK2 mutations and risk variants in Japanese patients with Parkinson’s disease. Mov Disord 24, 1034–1041 (2009).

78. Ross, O.A., et al. Analysis of Lrrk2 R1628P as a risk factor for Parkinson’s disease. Ann Neurol 64, 88–92 (2008).

79. Wang, X., Zhang, X., Xue, L. & Xie, A. The association between the LRRK2 R1628P variant and the risk of Parkinson’s disease in Asian: a meta-analysis. Neurosci Lett 623, 22–27 (2016).

80. Zhang, P., et al. Association of LRRK2 R1628P variant with Parkinson’s disease in Ethnic Han-Chinese and subgroup population. Sci Rep 6, 35171 (2016).

81. Chao, Y.X., et al. Lrrk2 R1628P variant is a risk factor for essential tremor. Sci Rep 5, 9029 (2015).

82. Zhao, Y., et al. LRRK2 variant associated with Alzheimer’s disease. NeurobiolAging 32, 1990–1993 (2011).

83. Ross, O.A., et al. Association of LRRK2 exonic variants with susceptibility to Parkinson’s disease: a case-control study. Lancet neurology 10, 898–908 (2011).

