## supplementary table 4 for "Pleiotropic effects for Parkin and LRRK2 in leprosy type-1 reactions and Parkinson’s disease"

| Table S3. Annotation of exonic variants identified in the T1R targeted genes. | | | | | | | | | | | | | |
| --- | --- | --- | --- | --- | --- | --- | --- | --- | --- | --- | --- | --- | --- |
| **Chr** | **Pos hg19** | **dbSNP** | **Cataloged** | **REF** | **ALT** | **Gene** | **Func** | **AA Change** | **SIFT score (pred)** | **Polyphen2 HDIV score (pred)** | **LRT score (pred)** | **MutationTaster score (pred)** | **CADD phred** |
| 4 | 38798399 | rs187624160 | dbSNP | G | T | *TLR1* | nonsynonymous | p.T685N | 1 (T) | 0 (B) | 0.026 (N) | 1 (N) | 0.002 |
| 4 | 38798417 | rs56205407 | dbSNP | A | G | *TLR1* | nonsynonymous | p.I679T | 0 (D) | 0.999 (D) | 0 (D) | 1 (D) | 26.4 |
| 4 | 38798648 | rs5743618 | dbSNP | C | A | *TLR1* | nonsynonymous | p.S602I | 1 (T) | 0 (B) | 0.224 (N) | 1 (P) | 2.562 |
| 4 | 38798694 | rs370009768 | dbSNP | C | T | *TLR1* | nonsynonymous | p.V587I | 0.627 (T) | 0.012 (B) | 0.905 (N) | 1 (N) | 0.699 |
| 4 | 38798755 | . | novel | T | C | *TLR1* | synonymous | p.L566L | . | . | . | . | . |
| 4 | 38798935 | rs5743614 | dbSNP | C | T | *TLR1* | synonymous | p.S506S | . | . | . | . | . |
| 4 | 38798971 | rs143576719 | dbSNP | T | C | *TLR1* | synonymous | p.V494V | . | . | . | . | . |
| 4 | 38799000 | . | novel | C | T | *TLR1* | nonsynonymous | p.G485R | 0.001 (D) | 0.866 (P) | 0.009 (N) | 0.991 (D) | 26.9 |
| 4 | 38799075 | rs137853171 | dbSNP | T | C | *TLR1* | nonsynonymous | p.I460V | 1 (T) | 0.043 (B) | 0.012 (N) | 0.997 (N) | 1.463 |
| 4 | 38799409 | rs200829133 | dbSNP | G | T | *TLR1* | nonsynonymous | p.S348R | 0.03 (D) | 0.938 (P) | 0 (D) | 1 (D) | 24.1 |
| 4 | 38799444 | rs200457447 | dbSNP | G | A | *TLR1* | nonsynonymous | p.R337C | 0.213 (T) | 0.004 (B) | 0.636 (N) | 1 (N) | 23 |
| 4 | 38799594 | . | novel | G | T | *TLR1* | nonsynonymous | p.L287M | 0.029 (D) | 0.833 (P) | 0.55 (N) | 1 (N) | 7.442 |
| 4 | 38799710 | rs4833095 | dbSNP | T | C | *TLR1* | nonsynonymous | p.N248S | 0.175 (T) | 0.001 (B) | 0.147 (N) | 1 (P) | 0.002 |
| 4 | 38800022 | rs117033348 | dbSNP | A | G | *TLR1* | nonsynonymous | p.L144P | 0 (D) | 1 (D) | 0 (D) | 1 (D) | 25 |
| 4 | 38800101 | rs5743612 | dbSNP | G | A | *TLR1* | nonsynonymous | p.H118Y | 0.141 (T) | 0.541 (P) | 0 (N) | 1 (P) | 1.459 |
| 4 | 38800323 | rs76600635 | dbSNP | A | G | *TLR1* | nonsynonymous | p.S44P | 1 (T) | 0 (B) | 0.172 (N) | 1 (N) | 0.001 |
| 4 | 38800339 | rs5743610 | dbSNP | G | A | *TLR1* | synonymous | p.H38H | . | . | . | . | . |
| 4 | 38802528 | rs5743596 | dbSNP | G | A | *TLR1* | UTR5 | . | . | . | . | . | . |
| 4 | 38805942 | rs5743566 | dbSNP | G | C | *TLR1* | UTR5 | . | . | . | . | . | . |
| 4 | 38805959 | rs145843781 | dbSNP | C | T | *TLR1* | UTR5 | . | . | . | . | . | . |
| 4 | 38805983 | rs5743565 | dbSNP | T | C | *TLR1* | UTR5 | . | . | . | . | . | . |
| 4 | 38806379 | rs1023321174 | dbSNP | A | G | *TLR1* | UTR5 | . | . | . | . | . | . |
| 4 | 38806384 | . | novel | C | G | *TLR1* | UTR5 | . | . | . | . | . | . |
| 4 | 154609176 | rs559003601 | dbSNP | A | C | *TLR2* | UTR5 | . | . | . | . | . | . |
| 4 | 154624115 | rs761297226 | dbSNP | A | T | *TLR2* | nonsynonymous | p.K19M | 0.088 (T) | 0.066 (B) | 0.592 (N) | 1 (N) | 7.06 |
| 4 | 154624238 | rs763540765 | dbSNP | CCAA | C | *TLR2* | nonframeshift_deletion | p.60_61del | . | . | . | . | . |
| 4 | 154624606 | . | gnomAD | G | A | *TLR2* | nonsynonymous | p.A183T | 0.004 (D) | 0.873 (P) | 0.005 (N) | 1 (N) | 25.5 |
| 4 | 154624656 | rs3804099 | dbSNP | T | C | *TLR2* | synonymous | p.N199N | . | . | . | . | . |
| 4 | 154624882 | rs538114024 | dbSNP | C | T | *TLR2* | stopgain | p.Q275X | . | . | 0.148 (N) | 1 (D) | 28.4 |
| 4 | 154624924 | rs780473983 | dbSNP | C | G | *TLR2* | nonsynonymous | p.L289V | 0.011 (D) | 0.911 (P) | 0.003 (N) | 1 (N) | 22.1 |
| 4 | 154625259 | rs144038898 | dbSNP | A | G | *TLR2* | synonymous | p.A400A | . | . | . | . | . |
| 4 | 154625409 | rs3804100 | dbSNP | T | C | *TLR2* | synonymous | p.S450S | . | . | . | . | . |
| 4 | 154625619 | rs200936495 | dbSNP | C | T | *TLR2* | synonymous | p.D520D | . | . | . | . | . |
| 4 | 154625775 | rs772226208 | dbSNP | C | A | *TLR2* | synonymous | p.G572G | . | . | . | . | . |
| 4 | 154625794 | rs751131041 | dbSNP | C | T | *TLR2* | nonsynonymous | p.R579C | 0.001 (D) | 0.972 (D) | 0.1 (N) | 1 (N) | 24.8 |
| 4 | 154625879 | . | novel | T | C | *TLR2* | nonsynonymous | p.V607A | 0.424 (T) | 0.001 (B) | 0.275 (N) | 1 (N) | 0.001 |
| 4 | 154626083 | . | novel | A | G | *TLR2* | nonsynonymous | p.H675R | 0 (D) | 1 (D) | 0 (D) | 1 (D) | 23.4 |
| 4 | 154626094 | rs190143186 | dbSNP | T | A | *TLR2* | nonsynonymous | p.F679I | 0.001 (D) | 0.994 (D) | 0 (D) | 1 (D) | 25.2 |
| 4 | 154626402 | rs5743709 | dbSNP | G | A | *TLR2* | synonymous | p.A781A | . | . | . | . | . |
| 4 | 154626441 | . | gnomAD | T | A | *TLR2* | UTR3 | . | . | . | . | . | . |
| 4 | 154626887 | . | novel | T | C | *TLR2* | UTR3 | . | . | . | . | . | . |
| 4 | 154627108 | rs534305860 | dbSNP | C | G | *TLR2* | UTR3 | . | . | . | . | . | . |
| 6 | 161769276 | . | novel | G | A | *PRKN* | UTR3 | . | . | . | . | . | . |
| 6 | 161769411 | rs1018991074 | dbSNP | C | T | *PRKN* | UTR3 | . | . | . | . | . | . |
| 6 | 161769437 | rs16892481 | dbSNP | G | A | *PRKN* | UTR3 | . | . | . | . | . | . |
| 6 | 161769555 | . | gnomAD | G | A | *PRKN* | UTR3 | . | . | . | . | . | . |
| 6 | 161769774 | rs191142442 | dbSNP | G | T | *PRKN* | UTR3 | . | . | . | . | . | . |
| 6 | 161769835 | rs3734464 | dbSNP | T | C | *PRKN* | UTR3 | . | . | . | . | . | . |
| 6 | 161770240 | rs113233227 | dbSNP | TTATC | T | *PRKN* | UTR3 | . | . | . | . | . | . |
| 6 | 161770252 | rs541658812 | dbSNP | T | G | *PRKN* | UTR3 | . | . | . | . | . | . |
| 6 | 161770456 | rs528792599 | dbSNP | C | T | *PRKN* | UTR3 | . | . | . | . | . | . |
| 6 | 161770479 | rs71653629 | dbSNP | G | A | *PRKN* | UTR3 | . | . | . | . | . | . |
| 6 | 161770503 | . | novel | G | A | *PRKN* | UTR3 | . | . | . | . | . | . |
| 6 | 161770508 | rs530999210 | dbSNP | A | G | *PRKN* | UTR3 | . | . | . | . | . | . |
| 6 | 161771116 | rs35125035 | dbSNP | G | T | *PRKN* | UTR3 | . | . | . | . | . | . |
| 6 | 161781182 | rs562362828 | dbSNP | T | C | *PRKN* | nonsynonymous | p.K408R | 0.183 (T) | 0.019 (B) | 0.019 (N) | 0.992 (N) | 23 |
| 6 | 161781219 | rs539917500 | dbSNP | T | C | *PRKN* | nonsynonymous | p.R396G | 0.378 (T) | 0.033 (B) | 0.003 (N) | 1 (N) | 7.681 |
| 6 | 161807855 | rs1801582 | dbSNP | C | G | *PRKN* | nonsynonymous | p.V380L | 0.607 (T) | 0 (B) | 0.831 (N) | 1 (P) | 3.433 |
| 6 | 161807897 | rs56092260 | dbSNP | G | A | *PRKN* | nonsynonymous | p.R366W | 0.001 (D) | 1 (D) | 0 (D) | 1 (D) | 34 |
| 6 | 161990424 | . | novel | T | C | *PRKN* | nonsynonymous | p.K299R | 0.173 (T) | 0.995 (D) | 0 (N) | 0.577 (D) | 18.14 |
| 6 | 161990432 | . | novel | G | C | *PRKN* | synonymous | p.S296S | . | . | . | . | . |
| 6 | 162206851 | rs769230602 | dbSNP | C | T | *PRKN* | nonsynonymous | p.R275Q | 0.017 (D) | 1 (D) | 0 (D) | 0.998 (D) | 35 |
| 6 | 162394341 | rs146173584 | dbSNP | C | T | *PRKN* | nonsynonymous | p.D243N | 0.041 (D) | 0.993 (D) | 0 (D) | 0.996 (N) | 25.2 |
| 6 | 162622197 | rs1801474 | dbSNP | C | T | *PRKN* | nonsynonymous | p.S167N | 0.229 (T) | 0.027 (B) | 0.665 (N) | 1 (P) | 14.32 |
| 6 | 162864389 | rs577232474 | dbSNP | G | A | *PRKN* | nonsynonymous | p.R42C | 0.069 (T) | 0.726 (P) | 0 (D) | 1 (D) | 23.4 |
| 9 | 117547060 | rs55958817 | dbSNP | G | A | *TNFSF15* | UTR3 | . | . | . | . | . | . |
| 9 | 117547378 | rs565538085 | dbSNP | C | T | *TNFSF15* | UTR3 | . | . | . | . | . | . |
| 9 | 117547489 | . | novel | C | T | *TNFSF15* | UTR3 | . | . | . | . | . | . |
| 9 | 117547772 | rs10114470 | dbSNP | T | C | *TNFSF15* | UTR3 | . | . | . | . | . | . |
| 9 | 117548105 | rs56305331 | dbSNP | C | T | *TNFSF15* | UTR3 | . | . | . | . | . | . |
| 9 | 117548752 | rs767719765 | dbSNP | C | T | *TNFSF15* | UTR3 | . | . | . | . | . | . |
| 9 | 117549739 | rs563544377 | dbSNP | A | C | *TNFSF15* | UTR3 | . | . | . | . | . | . |
| 9 | 117549891 | . | novel | C | A | *TNFSF15* | UTR3 | . | . | . | . | . | . |
| 9 | 117550363 | . | novel | GT | G | *TNFSF15* | UTR3 | . | . | . | . | . | . |
| 9 | 117550392 | rs1014042083 | dbSNP | T | C | *TNFSF15* | UTR3 | . | . | . | . | . | . |
| 9 | 117550673 | rs12686846 | dbSNP | G | A | *TNFSF15* | UTR3 | . | . | . | . | . | . |
| 9 | 117550809 | rs542552837 | dbSNP | C | T | *TNFSF15* | UTR3 | . | . | . | . | . | . |
| 9 | 117550966 | rs533359206 | dbSNP | G | A | *TNFSF15* | UTR3 | . | . | . | . | . | . |
| 9 | 117550975 | . | novel | T | A | *TNFSF15* | UTR3 | . | . | . | . | . | . |
| 9 | 117551158 | rs536175623 | dbSNP | G | A | *TNFSF15* | UTR3 | . | . | . | . | . | . |
| 9 | 117551453 | rs12682881 | dbSNP | C | T | *TNFSF15* | UTR3 | . | . | . | . | . | . |
| 9 | 117552072 | rs140202988 | dbSNP | G | C | *TNFSF15* | UTR3 | . | . | . | . | . | . |
| 9 | 117552191 | rs16931739 | dbSNP | T | C | *TNFSF15* | UTR3 | . | . | . | . | . | . |
| 9 | 117552241 | rs12682952 | dbSNP | C | T | *TNFSF15* | UTR3 | . | . | . | . | . | . |
| 9 | 117552801 | rs754459102 | dbSNP | G | A | *TNFSF15* | synonymous | p.N229N | . | . | . | . | . |
| 9 | 117552885 | rs3810936 | dbSNP | T | C | *TNFSF15* | synonymous | p.V201V | . | . | . | . | . |
| 9 | 117553158 | rs16931745 | dbSNP | A | T | *TNFSF15* | nonsynonymous | p.F110L | 1 (T) | 0 (B) | 0.934 (N) | 1 (N) | 0.896 |
| 9 | 117553178 | rs550636247 | dbSNP | G | T | *TNFSF15* | nonsynonymous | p.Q104K | 0.054 (T) | 0.998 (D) | 0 (D) | 1 (D) | 22.8 |
| 9 | 117568331 | . | novel | A | G | *TNFSF15* | UTR5 | . | . | . | . | . | . |
| 9 | 117655664 | rs535844130 | dbSNP | G | A | *TNFSF8* | UTR3 | . | . | . | . | . | . |
| 9 | 117655707 | rs10435870 | dbSNP | T | A | *TNFSF8* | UTR3 | . | . | . | . | . | . |
| 9 | 117655933 | rs540279151 | dbSNP | A | C | *TNFSF8* | UTR3 | . | . | . | . | . | . |
| 9 | 117656286 | rs192297306 | dbSNP | A | G | *TNFSF8* | UTR3 | . | . | . | . | . | . |
| 9 | 117663496 | . | novel | A | G | *TNFSF8* | UTR3 | . | . | . | . | . | . |
| 9 | 117663527 | rs554677031 | dbSNP | G | T | *TNFSF8* | UTR3 | . | . | . | . | . | . |
| 9 | 117663550 | rs3181202 | dbSNP | T | C | *TNFSF8* | UTR3 | . | . | . | . | . | . |
| 9 | 117663800 | rs3181201 | dbSNP | G | C | *TNFSF8* | UTR3 | . | . | . | . | . | . |
| 9 | 117663884 | rs3181200 | dbSNP | G | T | *TNFSF8* | UTR3 | . | . | . | . | . | . |
| 9 | 117664136 | . | novel | AT | A | *TNFSF8* | UTR3 | . | . | . | . | . | . |
| 9 | 117664172 | rs2974 | dbSNP | T | C | *TNFSF8* | UTR3 | . | . | . | . | . | . |
| 9 | 117664211 | rs2295800 | dbSNP | T | C | *TNFSF8* | UTR3 | . | . | . | . | . | . |
| 9 | 117664316 | rs376452088 | dbSNP | T | C | *TNFSF8* | UTR3 | . | . | . | . | . | . |
| 9 | 117664380 | rs77691228 | dbSNP | C | T | *TNFSF8* | UTR3 | . | . | . | . | . | . |
| 9 | 117664444 | rs2295801 | dbSNP | G | A | *TNFSF8* | UTR3 | . | . | . | . | . | . |
| 9 | 117664573 | rs117673429 | dbSNP | C | T | *TNFSF8* | UTR3 | . | . | . | . | . | . |
| 9 | 117664681 | rs548167795 | dbSNP | C | T | *TNFSF8* | UTR3 | . | . | . | . | . | . |
| 9 | 117665187 | rs3181374 | dbSNP | A | G | *TNFSF8* | UTR3 | . | . | . | . | . | . |
| 9 | 117665193 | rs76486122 | dbSNP | C | T | *TNFSF8* | UTR3 | . | . | . | . | . | . |
| 9 | 117665357 | rs116851933 | dbSNP | C | T | *TNFSF8* | UTR3 | . | . | . | . | . | . |
| 9 | 117665379 | rs1126711 | dbSNP | A | G | *TNFSF8* | UTR3 | . | . | . | . | . | . |
| 9 | 117665435 | rs3181372 | dbSNP | A | G | *TNFSF8* | UTR3 | . | . | . | . | . | . |
| 9 | 117665570 | rs3181371 | dbSNP | C | G | *TNFSF8* | UTR3 | . | . | . | . | . | . |
| 9 | 117665703 | rs201591439 | dbSNP | GGA | G | *TNFSF8* | UTR3 | . | . | . | . | . | . |
| 9 | 117665707 | rs193019133 | dbSNP | G | C | *TNFSF8* | UTR3 | . | . | . | . | . | . |
| 9 | 117665709 | rs117760632 | dbSNP | T | C | *TNFSF8* | UTR3 | . | . | . | . | . | . |
| 9 | 117665752 | rs3181370 | dbSNP | T | C | *TNFSF8* | UTR3 | . | . | . | . | . | . |
| 9 | 117665885 | rs76391989 | dbSNP | G | A | *TNFSF8* | UTR3 | . | . | . | . | . | . |
| 9 | 117665916 | . | novel | C | G | *TNFSF8* | UTR3 | . | . | . | . | . | . |
| 9 | 117665922 | rs530765966 | dbSNP | G | T | *TNFSF8* | UTR3 | . | . | . | . | . | . |
| 9 | 117665931 | rs3181368 | dbSNP | A | T | *TNFSF8* | UTR3 | . | . | . | . | . | . |
| 9 | 117666020 | . | novel | A | T | *TNFSF8* | UTR3 | . | . | . | . | . | . |
| 9 | 117666056 | rs533057360 | dbSNP | G | T | *TNFSF8* | UTR3 | . | . | . | . | . | . |
| 9 | 117668142 | rs3181195 | dbSNP | C | T | *TNFSF8* | synonymous | p.R92R | . | . | . | . | . |
| 9 | 117692512 | rs575875443 | dbSNP | C | T | *TNFSF8* | synonymous | p.A24A | . | . | . | . | . |
| 9 | 117692788 | rs61760050 | dbSNP | T | G | *TNFSF8* | UTR5 | . | . | . | . | . | . |
| 9 | 117692842 | rs1034038996 | dbSNP | C | T | *TNFSF8* | UTR5 | . | . | . | . | . | . |
| 12 | 40626185 | rs144883021 | dbSNP | A | G | *LRRK2* | nonsynonymous | p.Q116R | 0.482 (T) | 0.005 (B) | 0.01 (N) | 0.795 (N) | 11.28 |
| 12 | 40634404 | rs201332859 | dbSNP | T | C | *LRRK2* | nonsynonymous | p.S231P | 0.342 (T) | 0.003 (B) | 0 (D) | 0.766 (N) | 15.79 |
| 12 | 40637439 | rs752123862 | dbSNP | G | A | *LRRK2* | nonsynonymous | p.R265K | 1 (T) | 0 (B) | 0.189 (N) | 1 (N) | 5.567 |
| 12 | 40643725 | rs41286466 | dbSNP | G | T | *LRRK2* | synonymous | p.A312A | . | . | . | . | . |
| 12 | 40645272 | rs201261152 | dbSNP | C | T | *LRRK2* | synonymous | p.A369A | . | . | . | . | . |
| 12 | 40657700 | rs7308720 | dbSNP | C | G | *LRRK2* | nonsynonymous | p.N551K | 0.009 (D) | 1 (D) | 0 (D) | 0.032 (P) | 27.3 |
| 12 | 40668701 | rs281865042 | dbSNP | A | G | *LRRK2* | nonsynonymous | p.K616R | 0.159 (T) | 0.012 (B) | 0 (N) | 0.979 (N) | 14.85 |
| 12 | 40671928 | . | novel | A | G | *LRRK2* | synonymous | p.V702V | . | . | . | . | . |
| 12 | 40677699 | rs34410987 | dbSNP | C | T | *LRRK2* | nonsynonymous | p.P755L | 0.597 (T) | 0.647 (P) | 0 (D) | 1 (A) | 25.1 |
| 12 | 40677810 | . | gnomAD | G | A | *LRRK2* | nonsynonymous | p.R792K | 0.732 (T) | 0.001 (B) | 0.031 (N) | 1 (N) | 0.007 |
| 12 | 40677926 | . | novel | A | G | *LRRK2* | nonsynonymous | p.K831E | 0.106 (T) | 0.816 (P) | 0.023 (N) | 1 (N) | 17.51 |
| 12 | 40688695 | rs7966550 | dbSNP | T | C | *LRRK2* | synonymous | p.L953L | . | . | . | . | . |
| 12 | 40692930 | . | novel | T | A | *LRRK2* | nonsynonymous | p.C1123S | 0.644 (T) | 0.004 (B) | 0.032 (N) | 1 (N) | 0.118 |
| 12 | 40696649 | . | novel | T | A | *LRRK2* | nonsynonymous | p.F1185L | 0.753 (T) | 0.983 (D) | 0 (D) | 0.999 (D) | 21.7 |
| 12 | 40702471 | rs201453370 | dbSNP | C | A | *LRRK2* | nonsynonymous | p.L1388I | 0.127 (T) | 0.001 (B) | 0 (D) | 0.878 (D) | 14.05 |
| 12 | 40702924 | rs773386479 | dbSNP | T | C | *LRRK2* | synonymous | p.Y1402Y | . | . | . | . | . |
| 12 | 40702987 | rs11175964 | dbSNP | G | A | *LRRK2* | synonymous | p.K1423K | . | . | . | . | . |
| 12 | 40704252 | rs74681492 | dbSNP | C | T | *LRRK2* | nonsynonymous | p.P1446L | 0.008 (D) | 1 (D) | 0 (D) | 1 (D) | 34 |
| 12 | 40707848 | rs575429866 | dbSNP | G | A | *LRRK2* | synonymous | p.E1537E | . | . | . | . | . |
| 12 | 40713834 | rs1427263 | dbSNP | C | A | *LRRK2* | synonymous | p.G1624G | . | . | . | . | . |
| 12 | 40713845 | rs33949390 | dbSNP | G | C | *LRRK2* | nonsynonymous | p.R1628P | 0.04 (D) | 1 (D) | 0 (D) | 1 (D) | 27.8 |
| 12 | 40713873 | rs11176013 | dbSNP | A | G | *LRRK2* | synonymous | p.K1637K | . | . | . | . | . |
| 12 | 40713901 | rs11564148 | dbSNP | T | A | *LRRK2* | nonsynonymous | p.S1647T | 0.953 (T) | 0 (B) | 0.209 (N) | 0.096 (P) | 3.16 |
| 12 | 40716260 | rs10878371 | dbSNP | T | C | *LRRK2* | synonymous | p.G1819G | . | . | . | . | . |
| 12 | 40716268 | rs558440102 | dbSNP | A | G | *LRRK2* | nonsynonymous | p.H1822R | 0.763 (T) | 0.404 (B) | 0.001 (D) | 0.705 (D) | 12.21 |
| 12 | 40722186 | rs745904571 | dbSNP | A | T | *LRRK2* | nonsynonymous | p.Y1894F | 0.083 (T) | 0.987 (D) | 0 (D) | 1 (D) | 27.7 |
| 12 | 40740686 | rs33995883 | dbSNP | A | G | *LRRK2* | nonsynonymous | p.N2081D | 0.081 (T) | 0.392 (B) | 0 (D) | 0.987 (D) | 24.7 |
| 12 | 40742254 | rs10878405 | dbSNP | G | A | *LRRK2* | synonymous | p.E2108E | . | . | . | . | . |
| 12 | 40753116 | . | novel | T | C | *LRRK2* | synonymous | p.L2300L | . | . | . | . | . |
| 12 | 40757328 | rs34778348 | dbSNP | G | A | *LRRK2* | nonsynonymous | p.G2385R | 0.188 (T) | 0.074 (B) | 0 (D) | 1 (N) | 16.81 |
| 12 | 40757330 | rs33962975 | dbSNP | A | G | *LRRK2* | synonymous | p.G2385G | . | . | . | . | . |
| 12 | 40758652 | rs3761863 | dbSNP | T | C | *LRRK2* | nonsynonymous | p.M2397T | 0.466 (T) | 0 (B) | 0.043 (N) | 1 (P) | 0.005 |
| 12 | 40760808 | . | novel | G | GAAGC | *LRRK2* | frameshift_insertion | p.G2464fs | . | . | . | . | . |
| 12 | 40761663 | rs66737902 | dbSNP | T | C | *LRRK2* | UTR3 | . | . | . | . | . | . |
| 12 | 40761898 | rs551302864 | dbSNP | A | G | *LRRK2* | UTR3 | . | . | . | . | . | . |
| 12 | 40761930 | . | novel | T | A | *LRRK2* | UTR3 | . | . | . | . | . | . |
| 12 | 40761931 | rs10878441 | dbSNP | A | C | *LRRK2* | UTR3 | . | . | . | . | . | . |
| 12 | 40761951 | rs3886747 | dbSNP | C | T | *LRRK2* | UTR3 | . | . | . | . | . | . |
| 12 | 40761972 | rs200503610 | dbSNP | C | T | *LRRK2* | UTR3 | . | . | . | . | . | . |
| 12 | 40762032 | rs199535249 | dbSNP | G | A | *LRRK2* | UTR3 | . | . | . | . | . | . |
| 12 | 40762110 | rs373184816 | dbSNP | C | T | *LRRK2* | UTR3 | . | . | . | . | . | . |
| 12 | 40762229 | . | gnomAD | A | ACACAGAAA CTCTCTTTGT | *LRRK2* | UTR3 | . | . | . | . | . | . |
| 12 | 40762288 | rs886049365 | dbSNP | G | T | *LRRK2* | UTR3 | . | . | . | . | . | . |
| 12 | 40762303 | rs1365770 | dbSNP | G | C | *LRRK2* | UTR3 | . | . | . | . | . | . |
| 12 | 40762509 | rs182229935 | dbSNP | T | C | *LRRK2* | UTR3 | . | . | . | . | . | . |
| 12 | 40762996 | rs142051987 | dbSNP | C | G | *LRRK2* | UTR3 | . | . | . | . | . | . |
| 12 | 40763013 | rs12422278 | dbSNP | T | A | *LRRK2* | UTR3 | . | . | . | . | . | . |
| 12 | 40763075 | rs199493658 | dbSNP | T | A | *LRRK2* | UTR3 | . | . | . | . | . | . |
| 16 | 50733599 | rs187264529 | dbSNP | G | A | *NOD2* | nonsynonymous | p.V92I | 0.094 (T) | 0.428 (B) | 0.005 (N) | 0.998 (D) | 22.3 |
| 16 | 50733859 | rs2067085 | dbSNP | C | G | *NOD2* | synonymous | p.S178S | . | . | . | . | . |
| 16 | 50733864 | . | gnomAD | G | A | *NOD2* | nonsynonymous | p.R180K | 0.809 (T) | 0.061 (B) | 0.001 (D) | 0.855 (D) | 4.384 |
| 16 | 50741800 | rs149071116 | dbSNP | C | T | *NOD2* | nonsynonymous | p.A192V | 0.003 (D) | 0.886 (P) | 0.334 (N) | 1 (N) | 21.9 |
| 16 | 50741840 | . | novel | AT | A | *NOD2* | frameshift_deletion | p.L206fs | . | . | . | . | . |
| 16 | 50744624 | rs2066842 | dbSNP | C | T | *NOD2* | nonsynonymous | p.P268S | 0.45 (T) | 0.029 (B) | 0.217 (N) | 1 (P) | 0.001 |
| 16 | 50744635 | rs773516123 | dbSNP | G | A | *NOD2* | synonymous | p.K271K | . | . | . | . | . |
| 16 | 50744753 | rs104895427 | dbSNP | C | T | *NOD2* | nonsynonymous | p.R311W | 0 (D) | 1 (D) | 0.433 (N) | 1 (D) | 32 |
| 16 | 50744772 | . | novel | C | CT | *NOD2* | frameshift_insertion | p.A317fs | . | . | . | . | . |
| 16 | 50744988 | rs528926956 | dbSNP | C | T | *NOD2* | nonsynonymous | p.T389M | 0.004 (D) | 1 (D) | 0.093 (N) | 0.995 (D) | 25.6 |
| 16 | 50745199 | rs2066843 | dbSNP | C | T | *NOD2* | synonymous | p.R459R | . | . | . | . | . |
| 16 | 50745211 | rs200656015 | dbSNP | C | T | *NOD2* | synonymous | p.P463P | . | . | . | . | . |
| 16 | 50745233 | rs1078327 | dbSNP | C | T | *NOD2* | nonsynonymous | p.R471C | 0.143 (T) | 0.005 (B) | 0.544 (N) | 1 (N) | 11.12 |
| 16 | 50745583 | rs1861759 | dbSNP | T | G | *NOD2* | synonymous | p.R587R | . | . | . | . | . |
| 16 | 50745661 | rs149870902 | dbSNP | C | T | *NOD2* | synonymous | p.F613F | . | . | . | . | . |
| 16 | 50753909 | rs201035873 | dbSNP | C | A | *NOD2* | nonsynonymous | p.Q902K | 0.31 (T) | 0.033 (B) | 0.005 (N) | 1 (D) | 19.29 |
| 16 | 50759442 | rs104895463 | dbSNP | C | T | *NOD2* | synonymous | p.L975L | . | . | . | . | . |
| 16 | 50765740 | . | novel | G | A | *NOD2* | UTR3 | . | . | . | . | . | . |
| 16 | 50765872 | rs527525652 | dbSNP | A | G | *NOD2* | UTR3 | . | . | . | . | . | . |
| 16 | 50766127 | rs3135499 | dbSNP | A | C | *NOD2* | UTR3 | . | . | . | . | . | . |
| 16 | 50766201 | . | novel | C | G | *NOD2* | UTR3 | . | . | . | . | . | . |
| 16 | 50766338 | . | novel | C | A | *NOD2* | UTR3 | . | . | . | . | . | . |
| 16 | 50766607 | rs140643942 | dbSNP | C | A | *NOD2* | UTR3 | . | . | . | . | . | . |
| 16 | 50766668 | rs548686827 | dbSNP | C | A | *NOD2* | UTR3 | . | . | . | . | . | . |
| 16 | 50766886 | rs3135500 | dbSNP | G | A | *NOD2* | UTR3 | . | . | . | . | . | . |
